# Genetic determinants of aerial root traits that support biological nitrogen fixation in maize

**DOI:** 10.1101/2025.05.30.657053

**Authors:** Daniel Laspisa, Rafael Venado, Rubens Diogo, Jean-Michel Ané, Jason G. Wallace

**Author notes:** Dirscoll’s of Florida, Dover, Florida, USA. Corresponding author: Jason Wallace.

## Abstract

Modern agriculture depends on chemically synthesized nitrogen fertilizer, which ensures high yields but also can carry significant environmental and economic costs. Biological nitrogen fixation (BNF) already supplies nitrogen to legume crops and several avenues of research are underway to extend it to non-legume crops. In maize (*Zea mays*), aerial roots have been shown to contribute to BNF in some varieties, and both having many aerial roots and large aerial roots contributes to the fixation trait. However, much of the genetics controlling aerial root number and size is still unknown. Here we validate and quantify BNF in maize varieties from Southern Mexico under controlled conditions and evaluate a population of double haploids derived from the elite inbred PHZ51 crossed with these varieties. We find that most aerial root traits (root number, nodes with roots, root size) are reasonably heritable (h^2^ 0.5-0.75) and generally uncorrelated with each other. QTL mapping identifies 5 QTL each affecting nodes with aerial roots and aerial root number per node; in both cases all but 1 QTL show an increase from the landrace allele. We also identify 11 QTL for aerial root diameter, with most positive QTL coming from PHZ51. Between the two populations, only a few QTL overlap, indicating a presumably high diversity of genes affecting aerial root morphology in landrace populations. Combining the best QTL into elite material may provide a path toward meaningful levels of BNF for maize, and additional work is needed to determine how viable this approach will be in field settings.

## Introduction

Modern agriculture is heavily dependent on synthetic nitrogen fertilizer produced via the Haber-Bosch process. Although such fertilizers enable high and consistent yields, their use is also associated with many drawbacks, including high costs (Vos *et al*. 2025) (Boline) (USDA ERS 2024), extensive infrastructure requirements (Snapp *et al*. 2023), and significant environmental impacts including water pollution (Liu *et al*. 2024) eutrophication (Camargo and Alonso 2006), and greenhouse gas emissions (Pan *et al*. 2022). These challenges underscore the urgent need to reduce dependency on synthetic nitrogen inputs.

One promising avenue lies in harnessing biological nitrogen fixation (BNF) for cereal crops. Unlike legumes, most cereals do not make strong associations with BNF-capable bacteria, which limits their nitrogen-use efficiency (Guo *et al*. 2023). Recent studies have highlighted the potential of certain traditional maize varieties from southern Mexico to host nitrogen-fixing bacteria on their aerial roots (Van Deynze *et al*. 2018; Bennett *et al*. 2020; Pankievicz *et al*. 2022). These landraces produce carbohydrate-rich mucus (mucilage) on the aerial roots that foster nitrogen-fixing bacterial communities capable of supplying between 30-80% of the nitrogen needs for the plant (Van Deynze *et al*. 2018). Some key traits positively correlate with improved BNF, including the number of nodes bearing aerial roots (Van Deynze *et al*. 2018) and a high number and diameter of aerial roots (Pankievicz *et al*. 2022).

Aerial roots are a unique, but poorly understood set of organs (Sparks 2023) found only in the Andropogoneae (maize, sorghum, and sugarcane) and Paniceae (foxtail millet) tribes (Hostetler *et al*. 2021). Some quantitative trait loci (QTL) and genome-wide association (GWAS) studies have been carried out for aerial root-related traits (Liu *et al*. 2022; Sun *et al*. 2022; Singh *et al*. 2023) and investigating waterlogging tolerance (Guo *et al*. 2021). However, there is scarce research on supporting BNF in maize. Understanding the genetic mechanisms underlying these traits is essential for leveraging them in breeding programs. Enhancing nitrogen fixation in maize and other cereals could reduce reliance on synthetic fertilizers, benefiting both the environment and agricultural sustainability (Wen *et al*. 2021).

Here, we validate and quantify BNF in maize varieties from Southern Mexico under controlled conditions and evaluate a population of double haploids derived from the elite inbred PHZ51 crossed with these varieties. This study estimates the heritability of brace root traits, examines correlations among these traits, and identifies genomic regions associated with them. This work sets the foundation for pre-breeding efforts aimed at improving BNF in maize, paving the way for more sustainable agricultural practices.

## Results

### Quantification of nitrogen from BNF in landraces from Southern Mexico

Previous research has shown that landraces from southern Mexico can acquire nitrogen through interactions between their aerial root mucilage and associated microbial communities (Van Deynze *et al*. 2018). We examined the biological nitrogen fixation potential of some landraces, previously reported to produce multiple nodes with aerial roots and mucilage, from the same geographical region (Wilker *et al*. 2024). Biological nitrogen fixation and uptake are difficult to directly assess, so we used a suite of methods to confirm them in this system.

We first assessed the nitrogen-fixing potential of the landrace Oaxa524 (CIMMYT accession: CB017456) using an acetylene reduction assay (ARA). Mucilage from greenhouse-grown plants of this accession was collected and inoculated with nitrogen-fixing and non-fixing bacteria isolated from the rhizosphere of cereals and the mucilage of maize and sorghum. **(Supplementary Figure S1, Supplementary Table ST1).** All nitrogen-fixing strains exhibited elevated ARA activity, with *Klebsiella michiganensis*, *Klebsiella variicola*, and *Azotobacter vinelandii* showing the highest values. Notably, both *Klebsiella* strains were isolated from maize and sorghum mucilage (**Figure 1A**). These findings indicate that mucilage provides a favorable environment for diazotrophic bacteria, with certain strains thriving and demonstrating superior nitrogen-fixing performance compared to others.

**Figure 1.**
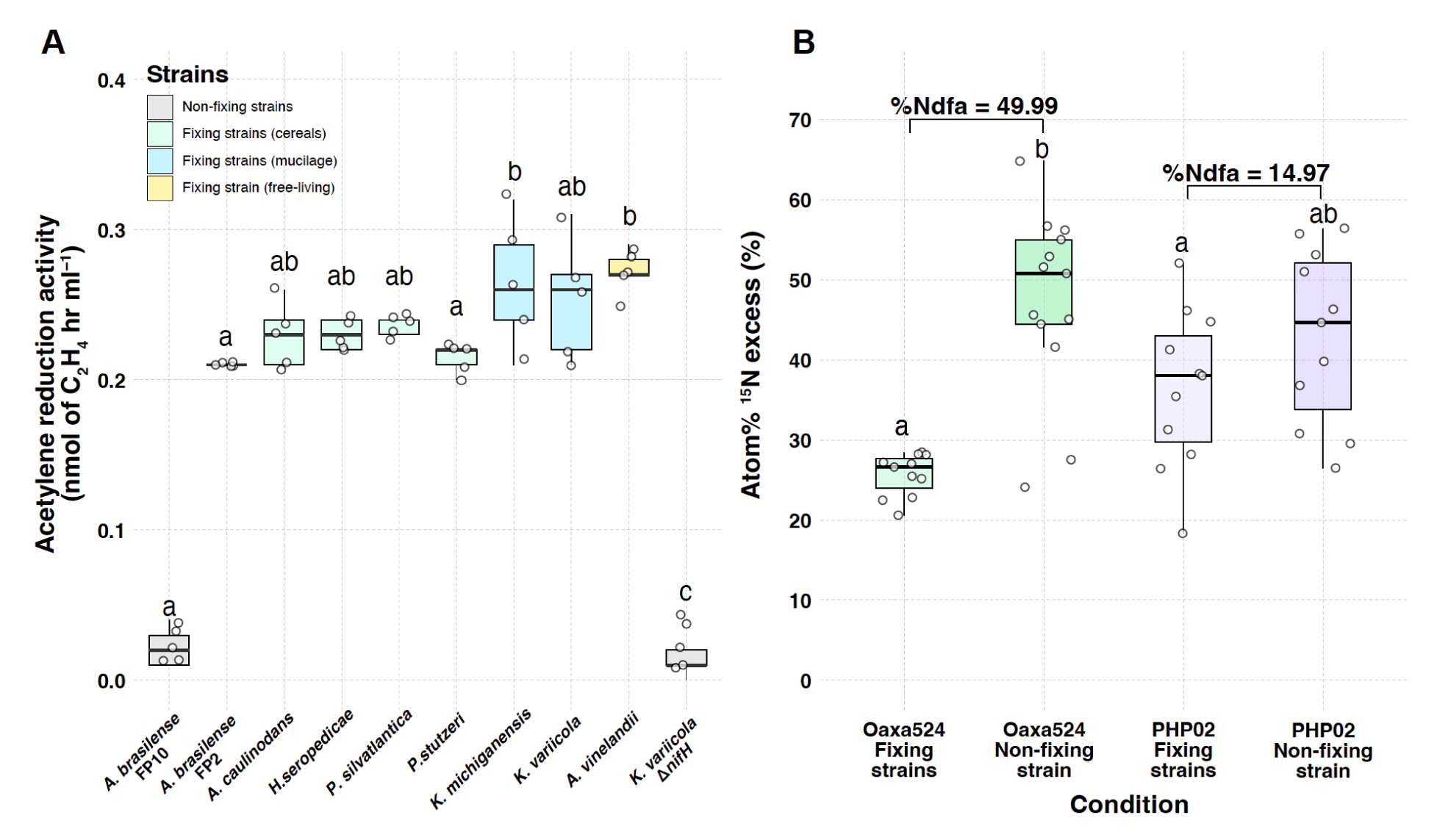
Assessment of biological nitrogen fixation in maize aerial roots. A) Acetylene reduction assay performed on maize mucilage grown in the greenhouse. The mucilage was used as a medium to cultivate ten strains: *Azospirillum brasilense* FP2, *Azorhizobium caulinodans*, *Azotobacter vinelandii* DJ100, *Herbaspirillum seropedicae*, *Klebsiella michiganensis*, *Klebsiella variicola*, *Paraburkholderia silvatlantica*, *Pseudomonas stutzeri* (nitrogen-fixing strains), and *Azospirillum brasilense* FP10 and *Klebsiella variicola* (non-nitrogen-fixing strains). Tukey’s honestly significant difference (HSD) test was used for statistical analysis in R, with significant groupings indicated in lowercase letters (n = 5). The four subcategories of strains are defined based on color codes and their specific roles or isolation. B) 15N isotope dilution experiment conducted on aerial roots of maize landrace Oaxa524 and exPVP PHP02. Both maize genotypes were inoculated with a mixture of *Klebsiella michiganensis* and *Azotobacter vinelandii* DJ100 (diazotrophic condition) or *Klebsiella michiganensis* ΔnifH (mutant strain). The entire plant at the flowering stage was ground for analysis of Isotope Ratio Mass Spectrometry (IRMS) and reported as Atom% ^15^N excess. Percent nitrogen derived from the atmosphere (Ndfa) was calculated for both maize genotypes, using plants inoculated with diazotrophs as the nitrogen-fixing condition and plants inoculated with mutant strains as the reference. ANOVA was performed to determine significant differences (n = 10-13), with the multcompView package used for analysis.

We performed a ^15^N isotope dilution assay to quantify the nitrogen uptake by Oaxa524 and compared it with the exPVP line PHP02, which produces negligible amounts of mucilage on its above ground roots. Both genotypes were grown in two separate greenhouses. In one greenhouse, plants were inoculated with nitrogen-fixing strains (*Azotobacter vinelandii* and *Klebsiella michiganensis*), selected for their high acetylene reduction activity (ARA), while in the other, plants were inoculated with a non-fixing mutant strain of *Klebsiella michiganensis* (**Supplementary Figure 1A**). At the flowering stage, we measured the Atom%^15^N excess using isotope ratio mass spectrometry (IRMS). For Oaxa524 inoculated with the fixing strains, the Atom%^15^N excess averaged 25%, whereas plants inoculated with the non-fixing strains showed higher Atom%^15^N excess values (**Figure 1B**). PHP02, whether inoculated with fixing or non-fixing strains, exhibited Atom%^15^N excess values similar to Oaxa524 inoculated with fixing strains. Although these results are ambiguous for PHP02, for Oaxa524 they clearly indicate plant uptake of fixed nitrogen from the applied bacteria.

To estimate nitrogen uptake from the atmosphere (%Ndfa) in these genotypes, we calculated %Ndfa at the flowering stage. Oaxa524 exhibited a %Ndfa of 49.99% (∼50%), while PHP02 recorded a %Ndfa of 14.97% (∼15%). Leaf samples collected earlier (at the V10 and V12-13 stages). Oaxa524 consistently showed low %Ndfa values for Oaxa524 (27-33%) and higher ones for PHP02 (17-33%), suggesting potential variability in nitrogen fixation efficiency across developmental stages.

### Development of doubled-haploid mapping populations

To better understand the genetics architecture of brace root traits we generated 8 populations of doubled haploids in collaboration with Limagrain, derived from an elite parent (PHZ51) crossed to one of three traditional varieties that exhibited traits conducive with improved symbioses with nitrogen fixing microbes. These traits include, aerial root diameter, aerial root number, and whirl number which are correlated with higher nitrogen fixation <u>(Pankievicz et al. 2022)</u>. We also measured stalk diameter as a covariate under the hypothesis that larger diameter stalks can accommodate more roots around the circumference. All 8 populations were genotyped however, due to logistical limitations we chose to focus field studies and phenotyping on the largest populations (Populations 1 and 7, derived from GRIN-PI-629238 and GRIN-PI-645940, respectively crossed to PHZ51).

The eight populations of doubled haploids were genotyped with an 1,800-marker SNP chip through Limagrain. A total of 1,574 DH lines were generated over 1,372 variable sites.

Population relatedness was visualized with multidimensional scaling (MDS) and to check for potential contamination in our materials (**Figure 2**). We do not observe any subclusters or individuals that would suggest genetic contamination in our populations (**Supplemental Figure S2**), and the distribution of minor allele frequencies were also in line with expected BC1 parameters (**Supplemental Figures S3**).

**Figure 2.**
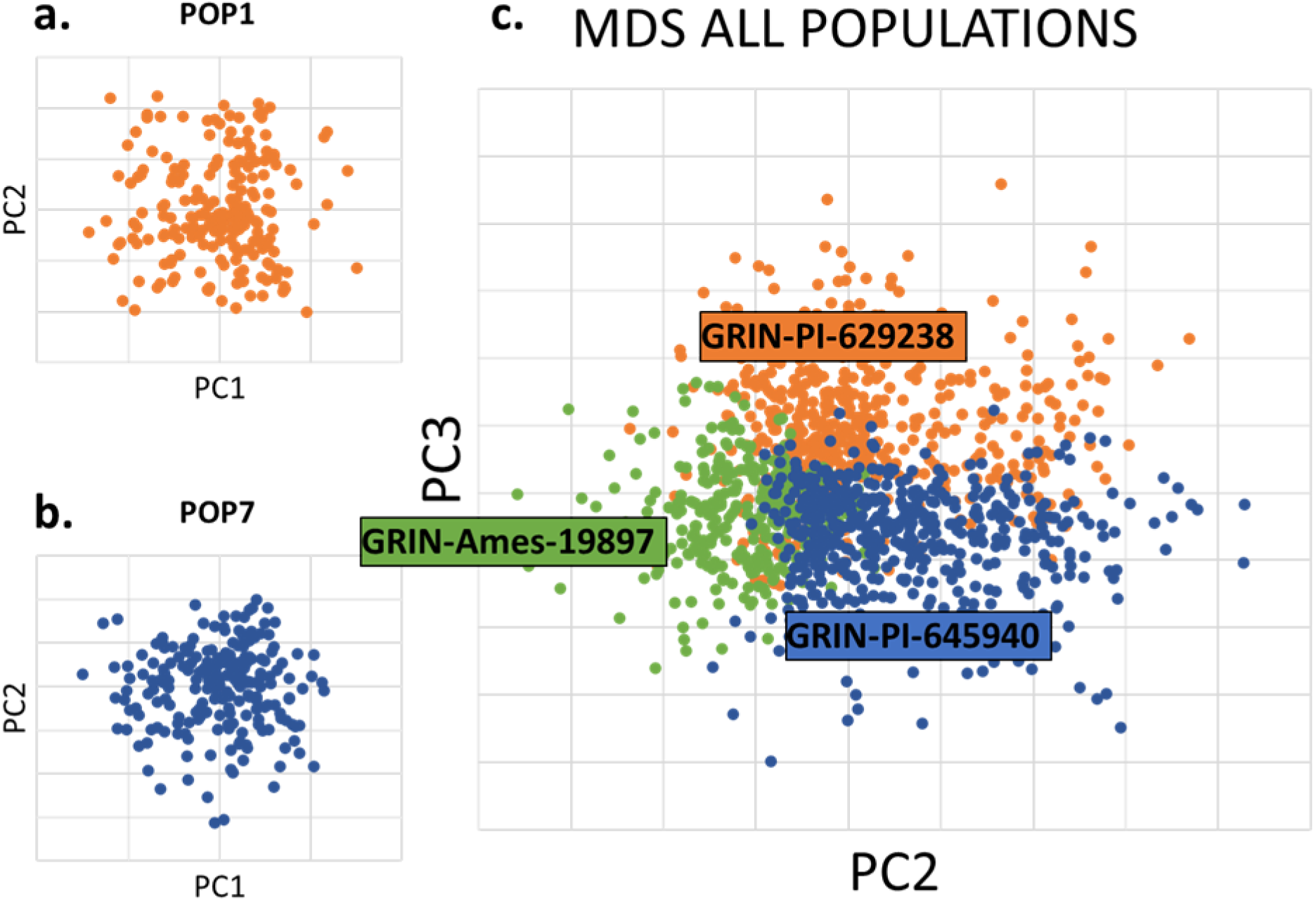
MDS plots of doubled haploids POP1 and POP7 and all 8 populations colored by landrace parent. We observe no secondary clusters or unexpected population structure within POP1 (Panel a.), or POP7 (panel b.). Panel c. comparing PC2 and PC3 shows a clear separation of the genotypes by landrace parent. PCI represents the genetic contribution of PHZ5l and is not shown. These MDS plots show no secondary clusters of genotypes and show the expected structure for 3 bi-parental crosses sharing a parent (PHZ5l). These distributions indicate there is likely no genetic contamination in our doubled haploid materials. MDS was performed in Tassel5.

Populations 1 and 7 were grown in Watkinsville, GA, and Hancock, WI, in 2023 and 2024 using a randomized complete block design with two replicates and 12 plants per plot. Brace root traits, including aerial root diameter, aerial root number, nodes with aerial roots, and stalk diameter, were measured after anthesis but before senescence to ensure we collected brace root data for mature plants. Measurements were taken on three plants per plot, with aerial root diameter assessed using a caliper, and root number counted on node four or the highest complete node. Nodes with aerial roots were recorded only on the main stem. Stalk diameter was measured on a single plant per plot across the narrowest part of the stalk and anthesis was recorded when 50% of a plot showed anther exertion on the main tassel spike.

### Distribution and heritability of traits

The distributions of five phenotypes (aerial root diameter, number, node number, stalk diameter and anthesis) were evaluated by separating the data by year and location (**Supplemental Figure S4**). We observe that the nodes with aerial roots phenotype were lower in Wisconsin in both 2023 and 2024 compared to Georgia. Aerial root number was higher in 2023 compared to 2024 across locations which may be attributed to environmental effects or a result of having different personnel doing the phenotyping in the Wisconsin location in 2024. Aerial root diameter and stalk diameter were observed to be higher in both locations in 2023 compared to 2024. Ultimately, these findings suggest brace root traits have considerable environmental variability.

Individual populations also showed variation in the traits (**Supplemental Figure S5**). Population 1 was observed to have more nodes with aerial roots than population 7. Population 7 was observed to have larger stalk diameters. Anthesis, aerial root number and diameter were relatively consistent across populations. Notably we observe that the PHZ51 elite parent has root diameters consistent with populations 1 and 7 whereas PH207 and PHP02 have smaller aerial root diameters.

Aerial root traits were generally normally distributed (**Supplemental Figure S6**). Heritability across aerial root traits was moderate with root diameter being the most heritable trait with an estimated 76% of phenotypic variation resulting from genetics (**Figure 3**). The heritability of nodes with aerial roots was estimated at 70% which is close to reported estimates of 89% (Li *et al*. 2024) (Li *et al*. 2024). The root number phenotype was the most environmentally dependent, with an estimated heritability of 59%. This is consistent with observations that plants at the ends of plots have higher numbers of aerial roots compared to those within the plots (edge effects) (**Supplemental Figure S7; Supplemental Text SX1; Supplemental Methods SM1**). The heritability of the control trait (anthesis, 56%) was considerably lower than reported estimates (Buckler *et al*. 2009) (Buckler *et al*. 2009), possibly because the backcross nature of these populations means they all flower relatively close to each other. Together these results suggest that aerial root traits are primarily controlled by genetics, indicating that meaningful progress can be made towards breeding for these traits.

**Figure 3.**
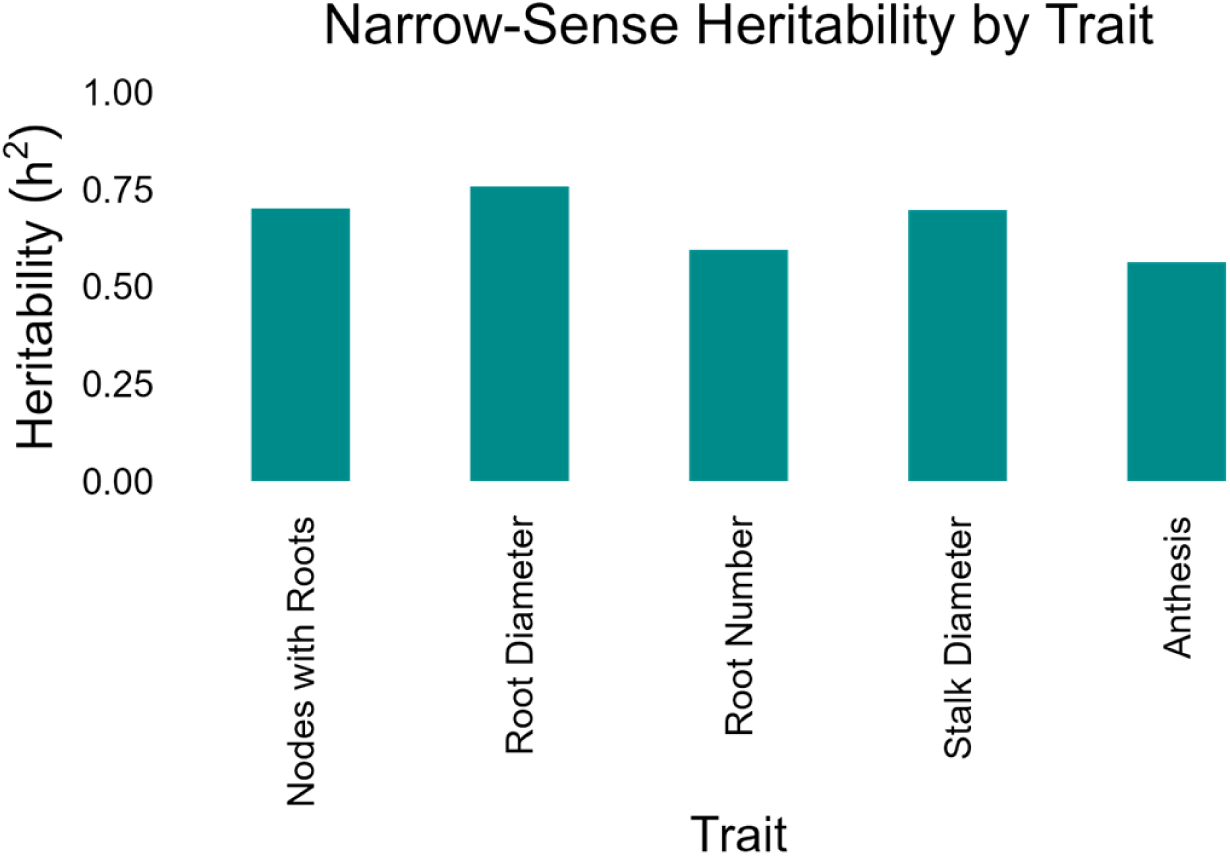
Narrow sense heritability plots for aerial root traits and anthesis for POP1 and POP7 combined phenotype data over 2023 and 2024.

### Correlations between aerial root traits

We conducted a pairwise correlation analysis to investigate correlations between aerial root traits (**Figure 4**). As expected, we observe moderate correlations between stalk diameter vs. root number (R^2^ ∼0.30) and stalk diameter vs. root diameter (R^2^ ∼0.22). Notably, we also observe a moderate positive correlation between aerial root number and nodes with aerial roots (R^2^ ∼0.41). While the correlation between root number and root diameter are statistically significant (p < 0.05), in practical terms the relationship is too weak to be meaningful (R^2^ ∼0.07). This contrasts with previous work that suggested the larger the roots, the fewer were present per whirl (Sun *et al*. 2022). The lack of correlation in our data may be due to a difference in genetics or a difference in phenotyping methodology (e.g., we counted roots on the top node, which is frequently less developed than lower nodes).

**Figure 4.**
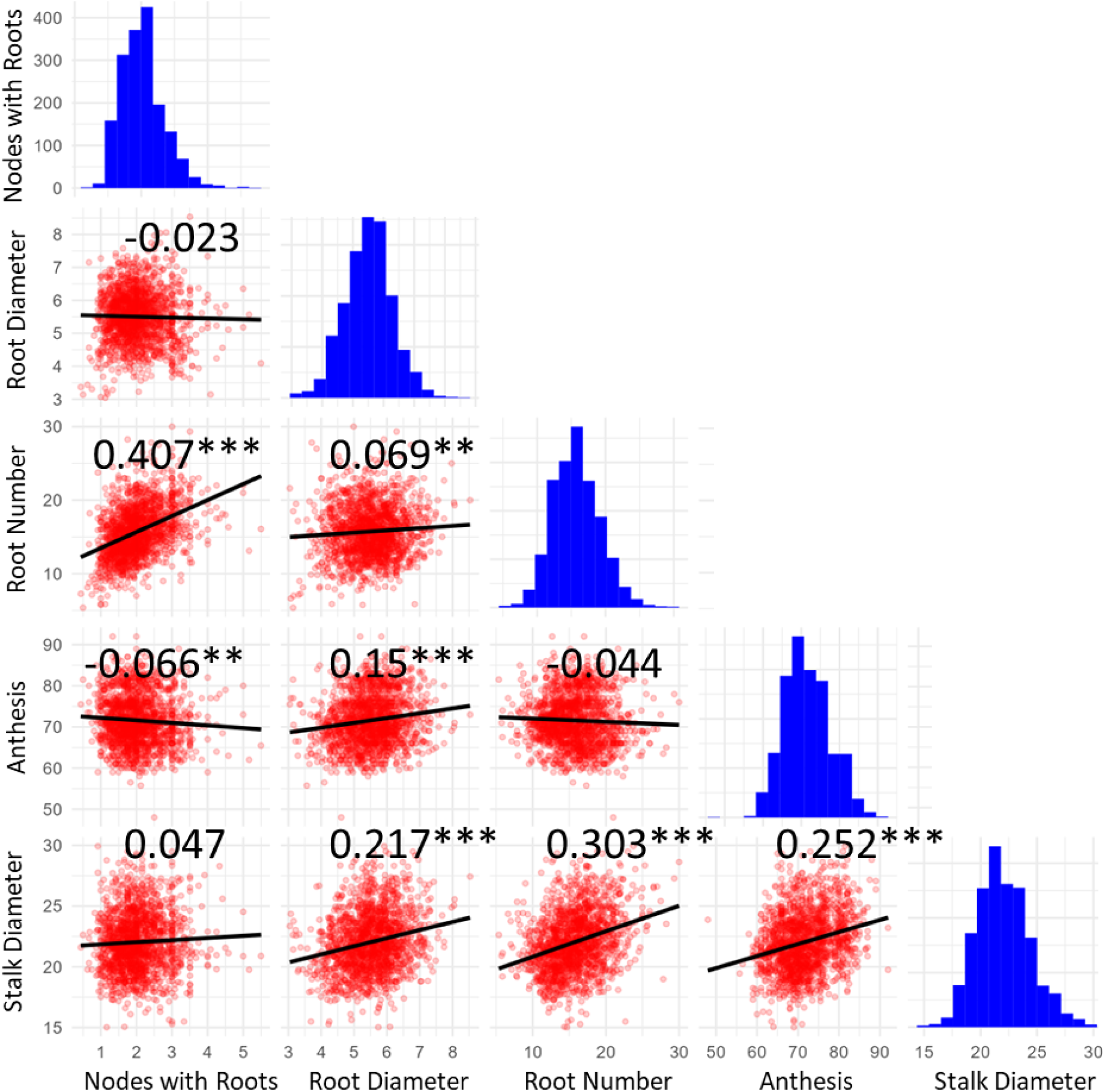
Pairwise correlation analysis for aerial root traits and the Anthesis control trait. The plots below the diagonal is a scatter plot of two traits. The diagonal plot represents the frequency of the phenotypic distribution for each of the traits. The value above the diagonal is Pearson's correlation coefficient between the two traits. * represents significant difference at the 0.05 level; ** represents significant differences at the 0.01 level.

### Mapping of aerial root traits

From these data, we performed quantitative trait locus (QTL) mapping to identify regions of the genome associated with brace root traits. Genotypes were filtered, removing non-mapped or non-segregating sites using Tassel5. Genotypes were imputed with FSFHap, and genetic maps for populations 1 and 7 were constructed with MSTmap and validated against the B73 v5 genome. Phenotypic data were processed in R, and BLUPs were calculated using lme4, accounting for location, year, and genotype. We estimated narrow-sense heritability with rrBLUP. QTL were identified using Haley-Knott regression in rqtl2, and parental effects were analyzed with rqtl2 in a separate analysis. The identified QTL were then compared to 409 published QTL/GWAS hits, with coordinates standardized to B73 v5 for validation.

We observe a total of 16 unique QTL including ten from population 1 and ten from population 7 (**Figure 5**), four of which appear to be shared. We observe three QTL for the aerial root number trait (one from POP1 and two from POP7) localizing to chromosomes 2, 8 and 10. There are nine QTL associated with the aerial root diameter trait (five from POP1 and four from POP7) localizing to chromosomes 1, 2, 3, 4, 7, 8 and 10. Four QTL for the nodes with aerial root trait localized to chromosomes 1, 3 and 9. Population 1 and population 7 share a QTL for this trait on chromosome 9. Notably the QTL on chromosome 9 is remarkably small in population 1 (only ∼8 Mb); most QTL are larger simply due to the relatively few recombinations involved in creating a BC1 population.

**Figure 5.**
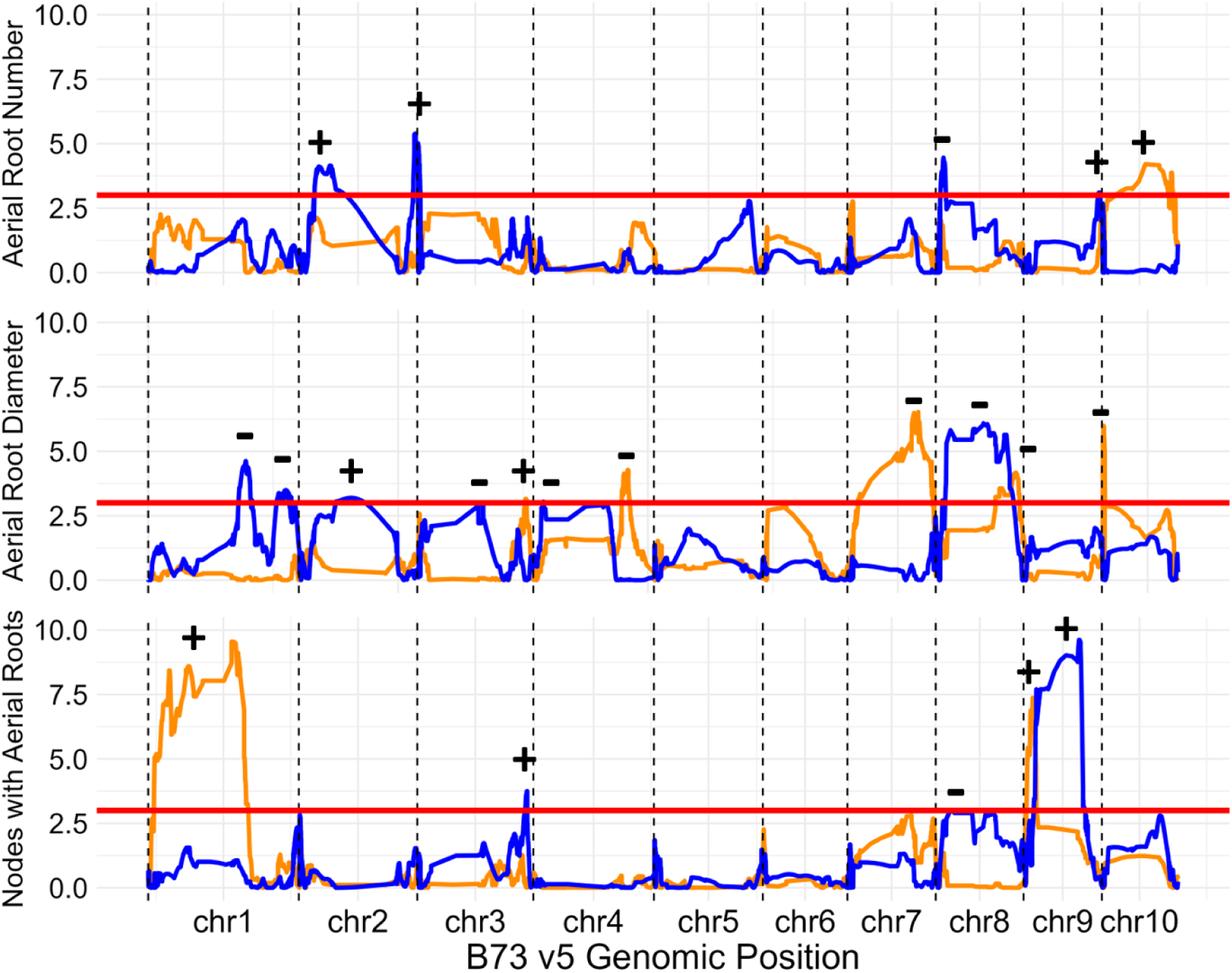
Genomic positions of QTLfor populations 1 (orange) and 7 (blue) for three aerial root traits. QTL positions are shown in the context of the B73 v5 genome. The y axis is the LOD score for each of the threetraits. Each QTL is marked with a + if the landrace parent has a positive effect on the trait and − if the landrace parent has a negative effect on the trait. The significance threshold for each trait is marked as a horizontal red line. The large QTL size is a result of limited recombination in our populations. Parental effects of each QTL can be found in **Supplemental Figure S8.**

### Comparison to published nodal root loci

We next compared these QTL to published root architecture loci (QTL and GWAS hits) from 10 studies (Burton *et al*. 2014, Chen *et al*. 2022, Gu *et al*. 2017, Ku *et al*. 2012, Liu *et al*. 2023, Moussa *et al*. 2021, Sun *et al*. 2020, Sun *et al*. 2022, Wang *et al*. 2019, Zhang *et al*. 2018) to look for potentially common genetic loci. We identified 339 published hits that overlap with our QTL (**Supplemental Figure S9**). Of those that overlap we find that 85 published GWAS hits overlap with our QTL and share the same trait (i.e. root number, root diameter, number of nodes, etc.). These 85 loci were compared to the annotated B73 v5 genome to identify putative candidate genes underlying these traits. 20 putative candidate genes were uncovered, including 17 for the aerial root number trait and 3 for the nodes with aerial roots trait (**Supplemental Table ST2**). We were unable to uncover putative candidate genes for the aerial root diameter trait using this method. These candidate genes will serve as the basis for future functional analysis validation of these results.

## Discussion

### Nitrogen fixation is supported in maize

In a previous study by our group, we evaluated the number of nodes with aerial roots in fixation-association landraces of maize and found that they developed nodes even during the adult stage, a feature absent in exPVP varieties (Wilker *et al*. 2024). A distinguishing characteristic of these landraces is their secretion of carbohydrate-rich mucilage, a trait proven essential for nitrogen fixation on these roots. This phenomenon has been documented not only in cereals such as maize (Van Deynze *et al*. 2018) and sorghum (Venado *et al*. 2023) but also in the tropical shrub *Heterotis rotundifolia* (Pang *et al*. 2023). The mucilage supports unique microbial communities that colonize and provide nitrogen to the plant. Using the acetylene reduction assay (ARA) with various microbial strains in maize, we observed that certain strains outperformed others, highlighting the selective nature of the mucilage (**Figure 1A**). This selective interaction was also confirmed in a sorghum study (Venado *et al*. 2023), where the same strain panel displayed varying levels of nitrogen fixation. Notably, when comparing results between the maize and sorghum studies, maize mucilage appeared to provide a more favorable environment for a broader range of diazotrophic bacteria (**Figure 1A**).

Sierra Mixe landraces have been reported to acquire between 29 and 89% of their nitrogen through symbiotic nitrogen fixation (Van Deynze *et al*. 2018). In our greenhouse experiments, we observed nitrogen acquisition at approximately 50% (**Figure 1B**), consistent with our previous findings. In a prior study on sorghum (Venado *et al*. 2023), we highlighted the challenges associated with accurately quantifying nitrogen uptake through these systems, acknowledging the limitations of various methodologies (Venado *et al*. 2023). Consequently, we anticipate variability in results under field conditions. Nonetheless, under controlled environments, this genotype and its derivatives represent a valuable genetic resource for breeding programs and provide a solid foundation for investigating the genetic regions regulating aerial root development.

### Prospects for nitrogen fixation in temperate maize

The findings described in this manuscript show that the development of nitrogen-fixing temperate maize holds promise. Reasonably high heritability estimates and the identification of aerial root trait associated genomic loci strongly suggest the trait can be selected for and improved in temperate maize. However there are many challenges and open questions that must be addressed. Specifically, water requirements, dependency on diazotrophic microbes, and potential yield reductions.

There are also some significant limitations to the trait namely water availability, and the presence of sufficient diazotrophic microbes. Mucilage production in aerial roots is dependent on the presence of water, usually rain. The minimum level of moisture required to stimulate mucilage production is as of yet unknown but observations suggest high humidity is required and overhead irrigation in the field is sufficient to induce the production of aerial root mucilage (data not shown).

As nitrogen fixation in maize relies on symbiotic relationships of diazotrophic microbes and the plant, the presence of these microbes in the mucilage is required for the fixation of nitrogen. These microbes are generally soil-borne and move up to the mucilage as an aerosol created from rainfall. However, to ensure that 1) the microbes are present on the plants and 2) ensure high levels of nitrogen fixation, Inoculation with diazotrophs will probably be necessary during the first years of using new accessions with the trait. Re-inoculation may also be necessary if diazotrophic microbial populations dwindle over time.

There also remains an open question regarding the potential for yield losses as a tradeoff of acquiring the nitrogen fixation trait. However current rough estimates in maize suggest a potential loss of 2-11% (Van Gelder *et al*. 2023). However, more precise future calculations will likely narrow this estimate.

The use of traditional varieties and more exotic germplasm will also be required to fully employ biological nitrogen fixation in maize. Studies in Sorghum suggest that aerial root traits associated with nitrogen fixation were far more prevalent in landraces than in the Sorghum association panel (elite lines) suggesting there was selection against aerial root traits <u>(Wolf et al. 2024)</u>. This is likely consistent with maize, which would suggest that aerial root mucilage traits have been counter selected in cereals. Thus, screening and integration of these traits from exotic germplasm is required to identify and move the trait into elite maize.

## Materials and Methods

### Acetylene reduction assay

Mucilage from the landrace Oaxa524 was collected from a greenhouse experiment wherein plants were cultivated as described in a previous study (Wilker *et al*. 2024). Nitrogenase activity was assessed in ten bacterial strains grown in the mucilage. These strains were categorized into nitrogen-fixing strains (*Azospirillum brasilense* FP2, *Azorhizobium caulinodans*, *Herbaspirillum seropedicae*, *Paraburkholderia silvatlantica*, *Pseudomonas stutzeri*, *Klebsiella michiganensis*, *Klebsiella variicola*, and *Azotobacter vinelandii*) and non-fixing strains (*Azospirillum brasilense* FP10 and *Klebsiella michiganensis* Δ*nifH*). Bacteria were cultured overnight in 3 mL of LB medium at 30 °C with shaking at 180 rpm. The cultures were centrifuged at 5000 g for 5 minutes, and the pellets were resuspended in 1 mL of sterile water. Next, 30 µL (OD600nm = 0.1) were inoculated into sterile vials containing 3 mL of mucilage. The vials were sealed and incubated for 24 hours at 30 °C. Subsequently, 1 mL of acetylene was injected into the vials after air replacement, followed by incubation for 3 days at 30 °C. Gas Chromatography-Flame Ionization Detection (GC-FID) was performed using a Shimadzu GC-2010 equipped with an HS20 autosampler. A 1 mL sample was injected, with both the injector and detector maintained at 200 °C, a nitrogen carrier gas flow rate of 86.6 mL/min, and a column temperature of 100 °C. Control vials containing acetylene were included to monitor ethylene traces. Nitrogenase activity was indirectly quantified based on ethylene production and expressed in nmol of ethylene produced per hour per mL of mucilage.

### 15N dilution experiments

In two greenhouse rooms at the Walnut Street Greenhouses, University of Wisconsin–Madison, maize landrace Oaxa524 and exPVP PHP02 were cultivated (Wilker *et al*. 2024). Each room contained 13 plants per genotype and was grown in 23 L pots filled with Pro-Mix LP15 medium (Quakertown, Pennsylvania). Greenhouse conditions were maintained at 28 °C during the day and 25 °C at night, with a 12-hour photoperiod (6:00 AM to 6:00 PM). Humidifiers (Smart Fog, Reno, Nevada) maintained an 80% humidity level to support mucilage production. Four weeks after germination, plants were fertilized weekly with 500 mL of Hoagland’s No.2 Basal Salt mixture without nitrogen (1.34 g/L) and supplemented with 80% ammonium-^15^N sulfate (^15^NH_4_)_2_SO_4_ as the nitrogen source. At six weeks post-germination, when aerial roots were visible, plants in one room were inoculated with a bacterial mixture (OD = 0.1 resuspended in PBS) containing *Klebsiella michiganensis* and *Azotobacter vinelandii*, while plants in the second room were inoculated with a *Klebsiella michiganensis* Δ*nifH* mutant. Each plant was fully sprayed with 250 mL of the inoculum suspended in phosphate-buffered saline (PBS) using a backpack sprayer. Samples were collected at specific growth stages: the third leaf (counting from the top) was harvested at the V10 stage and V12 post-inoculation. At flowering (VR1), entire plants were harvested, ground, and homogenized into a fine powder. The ^15^N content, measured as Atom% ^15^N excess above atmospheric levels, was analyzed via isotope ratio mass spectrometry (IRMS) in the Department of Soil Science, University of Wisconsin–Madison. The percentage of nitrogen derived from the atmosphere (%Ndfa) was calculated using the formula described in van Deynze et al. (2018). Plants inoculated with diazotrophs were considered “fixers,” while those inoculated with the mutant strain served as the reference group.

### Population development and genotyping

Eight populations of doubled haploid maize lines were developed in collaboration with Limagrain. The process began with haploid induction, where F1 plants, derived from crosses between PHZ51 and an exotic line, were pollinated with a male haploid inducer genotype (**Supplemental Figure S10**). Haploids were identified at the seed stage, and haploid (D0) seedlings were treated with colchicine to double their chromosome number. Fertile D0 plants were subsequently selfed to produce the final doubled haploid populations. These populations were genotyped using a chip with 1,800 markers, yielding 1,574 DH lines across 1,372 variable sites.

### Field experiments and phenotyping

Populations 1 and 7 were grown in Watkinsville, Georgia (33°43’25.9“N 83°18’19.8”W) and Hancock Wisconsin (123°25E, 41°48N) in 2023 and 2024 in a randomized complete block design with two replicates and 12 plants per plot. Plots measured 9 feet by 2.5 feet with 6-foot alleys separating ranges of plots. Brace root measurements were collected after anthesis but before senescence for aerial root diameter, aerial root number, the number of nodes with aerial roots, stalk diameter (**Figure 6**). <u>Aerial root diameter</u> was measured on three roots on three plants per plot using a digital caliper on three non-adjacent aerial roots from node four (or the highest node if the plant did not have four nodes with aerial roots) approximately 1-2cm from the stalk. <u>Aerial root number</u> was measured on three plants per plot, counting on node four. (If the plant did not have four nodes the number of roots on the highest complete node with roots was counted.) A complete whorl was defined as one with roots extending from the stalk at least halfway around the whorl. If the top-most node with roots was incomplete, we counted the number of roots in the previous node(s) until we reached a node with a complete whorl. Bumps <1cm were not counted as aerial roots. The <u>number of nodes with aerial roots</u> was measured on three plants per plot only on the main stem; tillers were not counted. Beginning at the soil level, we counted the number of nodes with visible roots, including the nodes with roots that touch the soil. Nodes were included if the roots extend at least 1 cm from the stalk. A rating of 0.5 was given when bumps (<1cm roots) were developing on the next node. For example, if a plant has six nodes with aerial roots longer than 1 cm and node 7 has root bumps, the number of nodes was recorded as 6.5. <u>Stalk diameter</u> was measured on a single representative plant from each plot after anthesis. The measurement was taken with a caliper across the narrower width of the elliptically shaped stalk, between nodes three and four. <u>Anthesis</u> was recorded as a date when 50% of a plot exhibited anther exertion on more than half of the main tassel spike.

**Figure 6.**
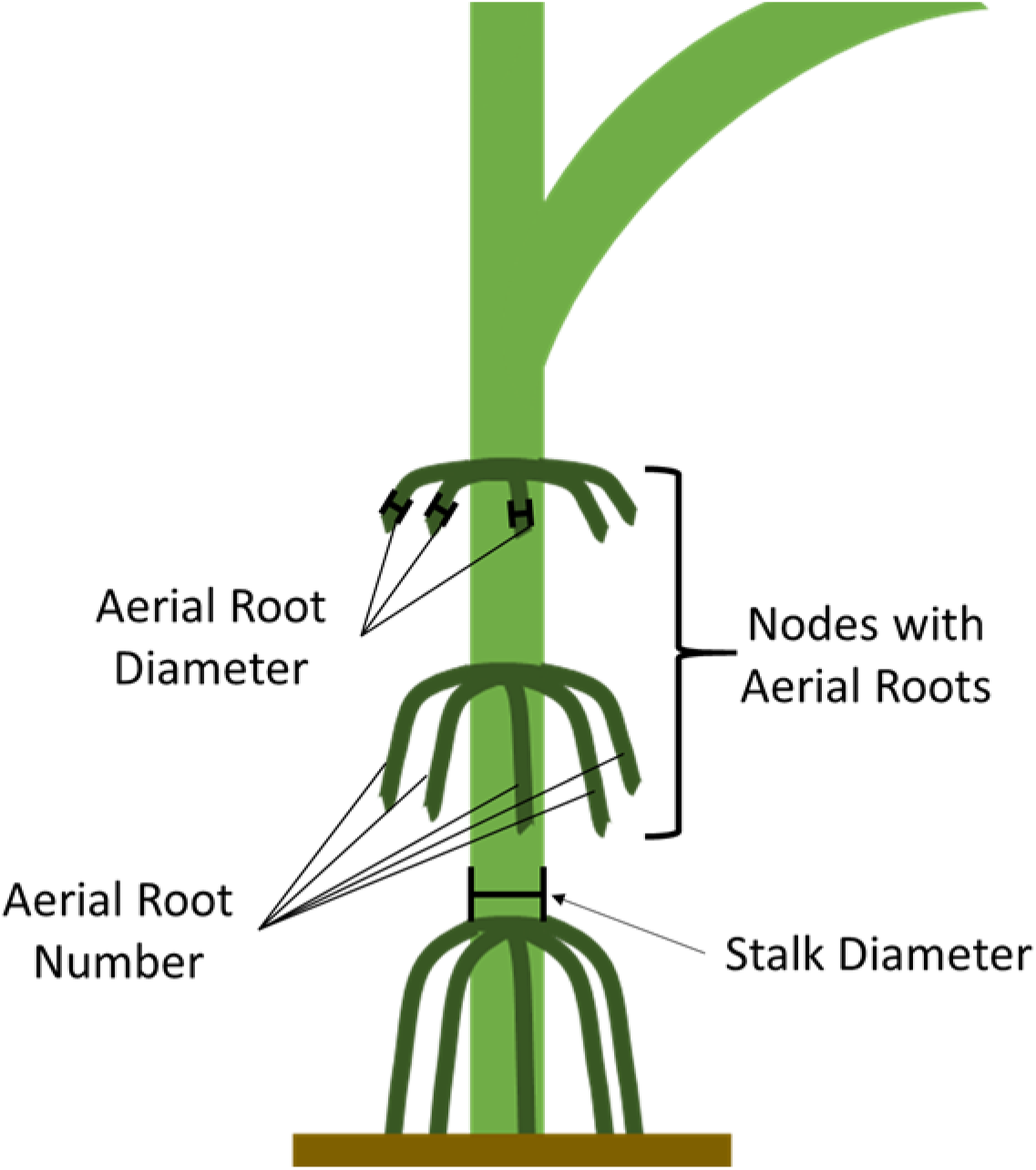
Schematic representation of the aerial root phenotypes collected in 2023 and 2024. Aerial root diameter, number, and the number of nodes with aerial roots have been correlated with increased mucilage production. Stalk diameter was collected as a covariate as a larger diameter stalk can accommodate more roots around its circumference.

### Genetic analysis

Initial genotype filtering removed 206 sites that had no corresponding coordinate position in the B73 v5 genome and/or were not segregating in populations 1 and 7. Genotypes were filtered in Tassel5 (Bradbury *et al*. 2007) allowing a maximum of 70% missing data, 7% max heterozygosity per site, a minimum allele frequency of 0.1. Genotypes for populations 1 and 7 were imputed with the Tassel5 FSFHap command line plugin (Swarts *et al*. 2014). Genetic maps were constructed independently with the filtered genotypes for populations 1 and 7 using MSTmap (Wu *et al*. 2008). Genetic maps were validated by comparing the positions of the genetic map to the physical map (B73 v5, see **Supplemental Figures S11 and S12**).

Phenotypes were processed in R. The distribution of phenotypes was plotted and visually inspected to identify outliers resulting from data entry errors in the field. Phenotypes were filtered in R to remove erroneous outliers with the following parameters: nodes with aerial roots (>=0, <=6), aerial root diameter (>=3, <=10), aerial root number (>=5, <=30), Stalk Diameter (>=15, <= 30), and Anthesis (>=40, <= 100). Post-filtering phenotype distributions were plotted to validate removal (**Supplemental Figures S5**). The filtered phenotypes were converted to best linear unbiased predictors (BLUPs) using a random effect model with the lme4 package in R (Bates *et al*. 2015) accounting for growth location, field year, and genotype. Narrow sense heritability estimates for brace root traits were calculated in R with the rrBLUP package (Endelman 2011).

Quantitative Trait Loci (QTL) were identified with Haley-Knott regression with the rqtl2 package in R (Broman *et al*. 2019). QTL were identified independently for populations 1 and 7. Parental effects for the identified QTL were identified with rql2.

QTL observed in our populations were compared to 409 QTL/GWAS hits from 10 publications associated with both maize root traits and “control” traits such as anthesis (see supplemental data and supplemental references). Coordinates of published QTL and GWAS hits were converted to B73 v5 physical map coordinates (Hufford *et al*. 2021) to validate overlap.

## Acknowledgements

We graciously thank Fabienne Henriot and Marie-Hélène Tixier of Limagrain for their contributions to the development of the doubled haploid populations used in this study. We thank Valentina Infante for generating the *Klebsiella michiganensis* Δ*nifH.* This work was funded by USDA-NIFA Grant 2020-08318.

## Table of Contents

### Supplemental Text

#### SX1. Aerial Root Traits are Impacted by Plot Position

Of the 2220 seeds sown, 1745 resulted in adult plants considered normal. One of the plots (J-78) had only one adult plant and was therefore excluded from the analyses since it would not be possible to estimate field edge effects in this way. In the remaining 110 plots, the 1744 plants were phenotyped for the five aerial root-related traits once mentioned. Results were normalized to a 0-1 range based on values presented by plants of the same genotype in the same row. In all five traits evaluated, we noted statistically significant differences according to the position of the plants within the row.

The number of nodes with aerial roots, the number of aerial roots at the top node, and the diameter of the plant stalk exhibited a U-quadratic-like distribution so that the values observed at the edges (deca-percentile groups 1 and 10 regarding plants’ position) were the highest, followed by groups 2 and 9 (i.e., those located adjacent to the rows’ ends); in contrast, the other groups (3 to 8), which constituted the most central portion of the row, presented the lowest results.

A similar situation occurred with the diameter of aerial roots at the top node; however, in this case, it was only possible to differentiate the results of the edge groups (1 and 10) from the others. In these four cases, the variance analyses indicated a p-value lower than 0.1%. Finally, the flag leaf height, one of the most heritable traits in maize, estimated to be higher than 90% (Peiffer et al. 2014), showed slightly more homogeneous results; even so, under a p-value ≤ 1%, the edge groups (1 and 10) exhibited a lower value than most of the other groups, except groups 4 and 7.

Based on these results, we conclude that the five aerial root-related traits evaluated here are influenced by the position of the plants in the experimental plots, given that the so-called field edge effects consistently affect the values observed for each of them.

## Supplemental Methods

### SM1. Estimation of Field Edge Effects

The trial was conducted at the West Madison Agricultural Research Station (43°03’56.0” N, 89°32’34.0” W) in Verona-WI, USA. The field management followed the agronomic practices for maize farming adopted in that region. In addition to the borders, one hundred eleven 12.5 ft (≈3.8 m)-long plots, spaced 2.5 ft (≈0.76 m) apart, were arranged. In each plot, twenty seeds of a given maize genotype were sown at a depth of 2″ (≈5 cm) and spaced 8″ (≈20 cm) apart. Forty-one genotypes were evaluated, with at least two replicates (plots) for each. The genotypes tested included three landraces with high numbers of aerial roots and late cycle, three inbred lines with expired Plant Variety Protection Act certificates (Ex-PVPs) adapted to the American Midwest region, and 35 lines originated from crosses between them, which are in advanced generations of back- or self-crossing and are part of a breeding program that aims to introgress aerial roots-related traits into those Midwest-adapted Ex-PVPs.

Excluding non-germinated seeds and plants that did not exhibit normal phenological development, all individuals from the 111 plots were phenotyped for the following traits: number of nodes with aerial roots, number of aerial roots at the top node, diameter of aerial roots at the top node (in mm), diameter of plant stalk (in mm), and flag leaf height (in cm). To facilitate the interpretation of results, they were normalized on a range from 0 to 1 using the following formula: normalized value = (value observed in that plant ‒ lowest value observed in the plot) / (highest value observed in the plot ‒ lowest value observed in the plot).

The position that each plant occupied in the row (1 to 20 in the plots in which all the plants appropriately developed but variable in the other plots) was also noted and normalized using the following formula: normalized position = (position of that plant within the row ‒ position of the first plant within the row) / (position of the last plant within the row ‒ position of the first plant within the row). Finally, we could classify the plants into ten deca-percentile groups based on this normalized position. Groups 1 and 10 constituted the row extremities; therefore, plants in these groups were the most exposed to edge effects.

## Supplemental Figures

**Figure S1.**
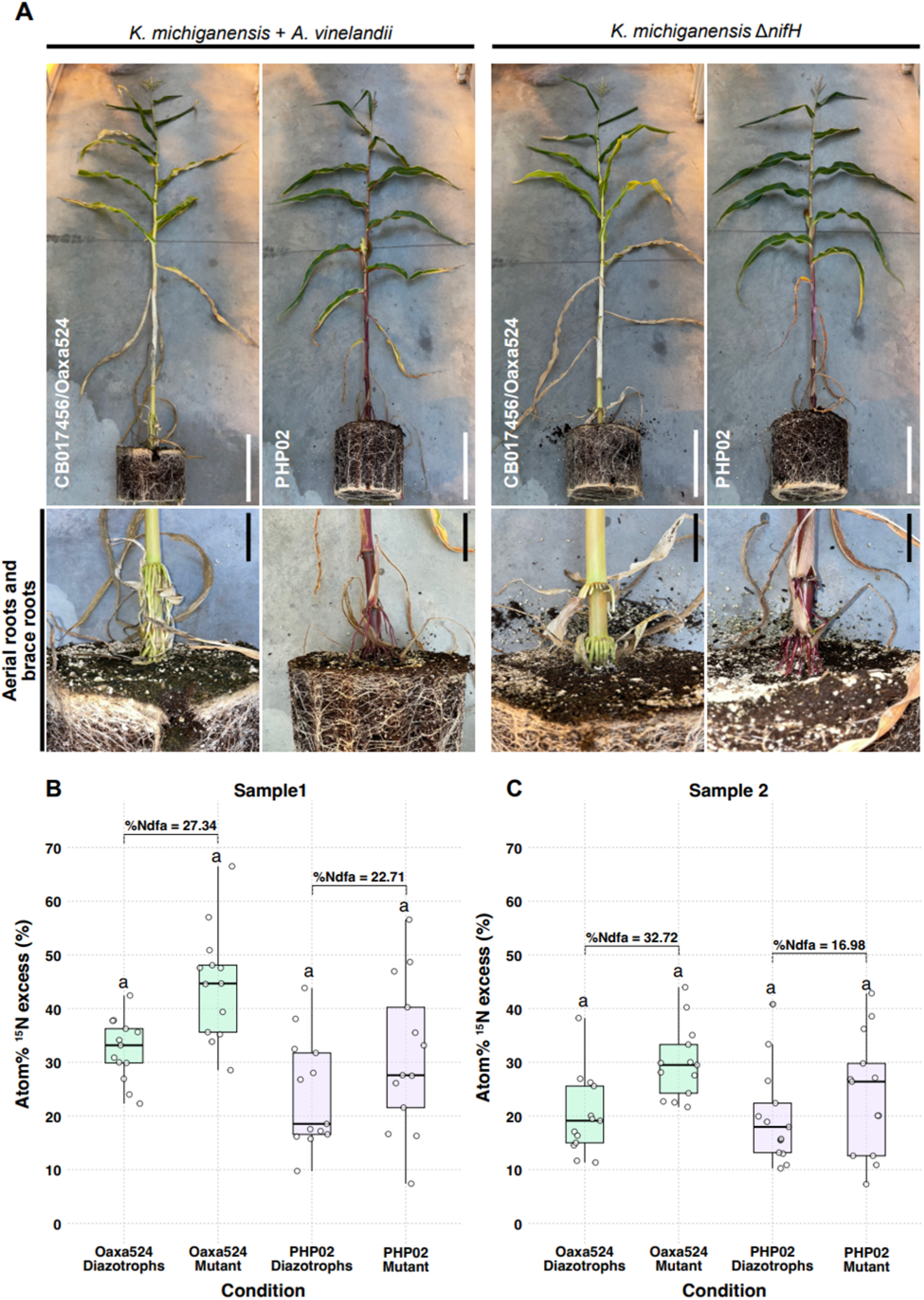
Determination of nitrogen fixation in maize. A) Flowering stage maize genotypes Oaxa524 and PHP02 were inoculated with: i) nitrogen-fixing diazotrophs (*K. michiganensis* and *A. vinelandii*) and ii) non-nitrogen-fixing diazotrophs (*K. michiganensis* ΔnifH). Scale bar = 25 cm. The lower panel shows the presence of aerial roots (Oaxa524) or brace roots (PHP02) under the two inocula. Scale bar = 8 cm B) At the V10 stage, leaf samples were collected to measure 15N isotope levels using IRMS. C) At the V12-13 stage, leaf samples from genotypes Oaxa524 and exPVP PHP02 were collected for 15N isotope analysis by IRMS. The genotypes were inoculated with either a mixture of *Klebsiella michiganensis* and *Azotobacter vinelandii* DJ100 (diazotrophic condition) or *Klebsiella michiganensis* ΔnifH (mutant strain). The percentage of nitrogen derived from the atmosphere (Ndfa) was calculated for both maize accessions, using diazotroph-inoculated plants as the nitrogen-fixing reference and mutant strain-inoculated plants as the non-fixing reference. ANOVA was conducted to identify significant differences, with the multcompView package utilized for the analysis (n = 10-13).

**Figure S2.**
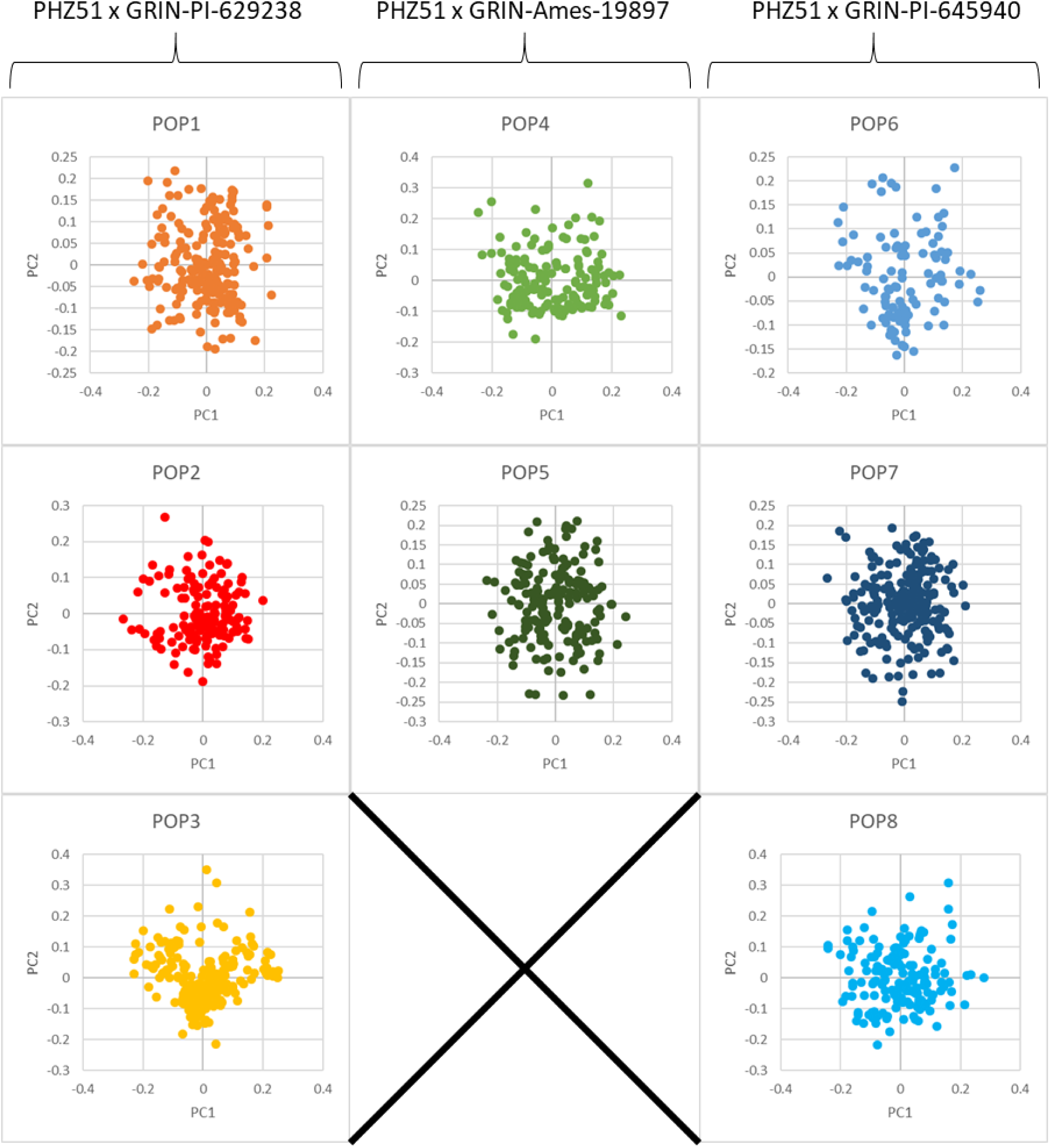
Multidimensional Scaling plots separated by population. MDS plots for individual populations show no secondary clusters of genotypes nor any clear sub-population structure indicating there is likely no genetic contamination in our doubled haploid materials. MDS was performed in Tassel5.

**Figure S3.**
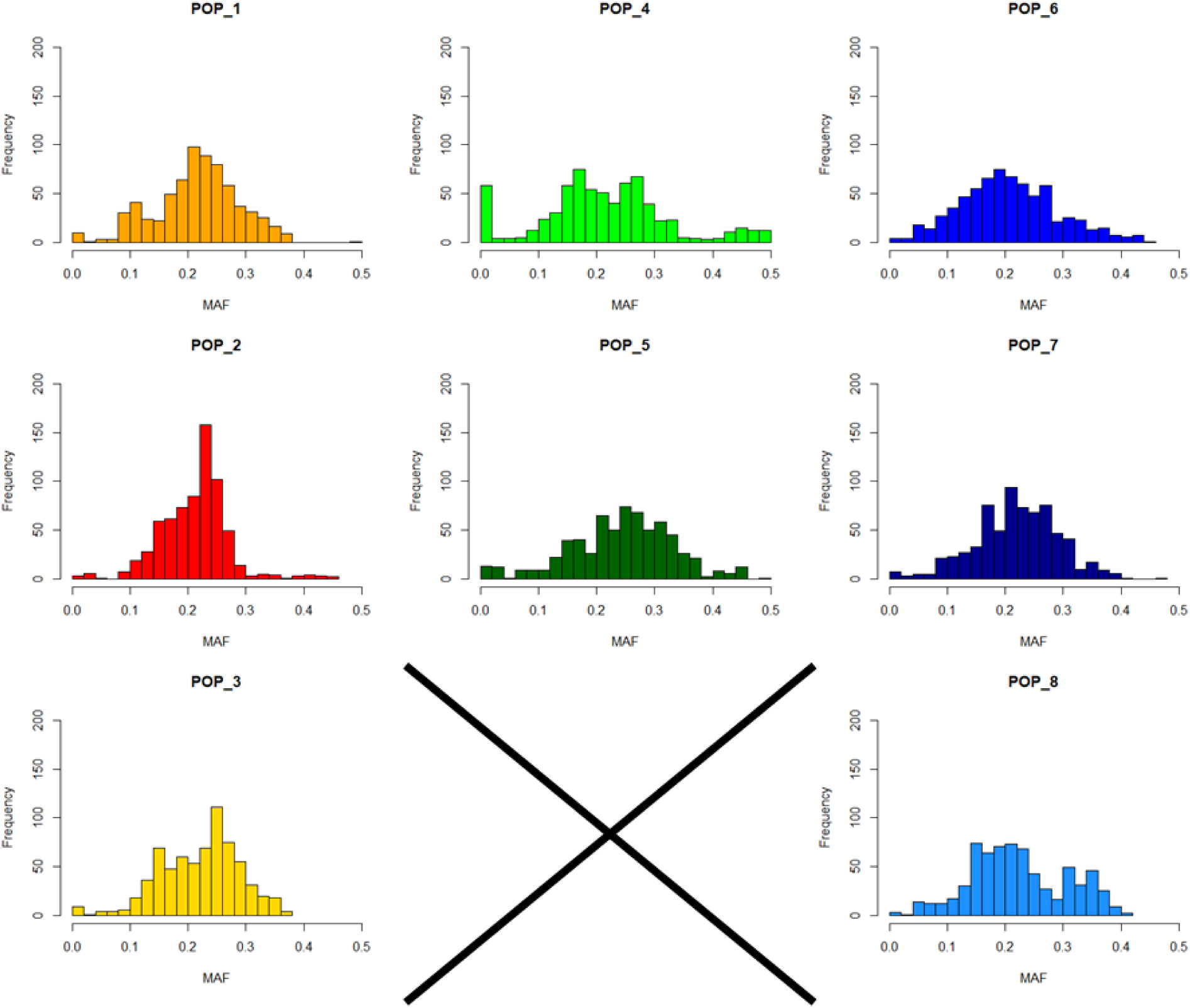
Minor Allele Frequency plots separated by population. Minor allele frequencies for all populations are roughly normally distributed and centered on ∼0.25 which is within expectations for our crossing scheme.

**Figure S4.**
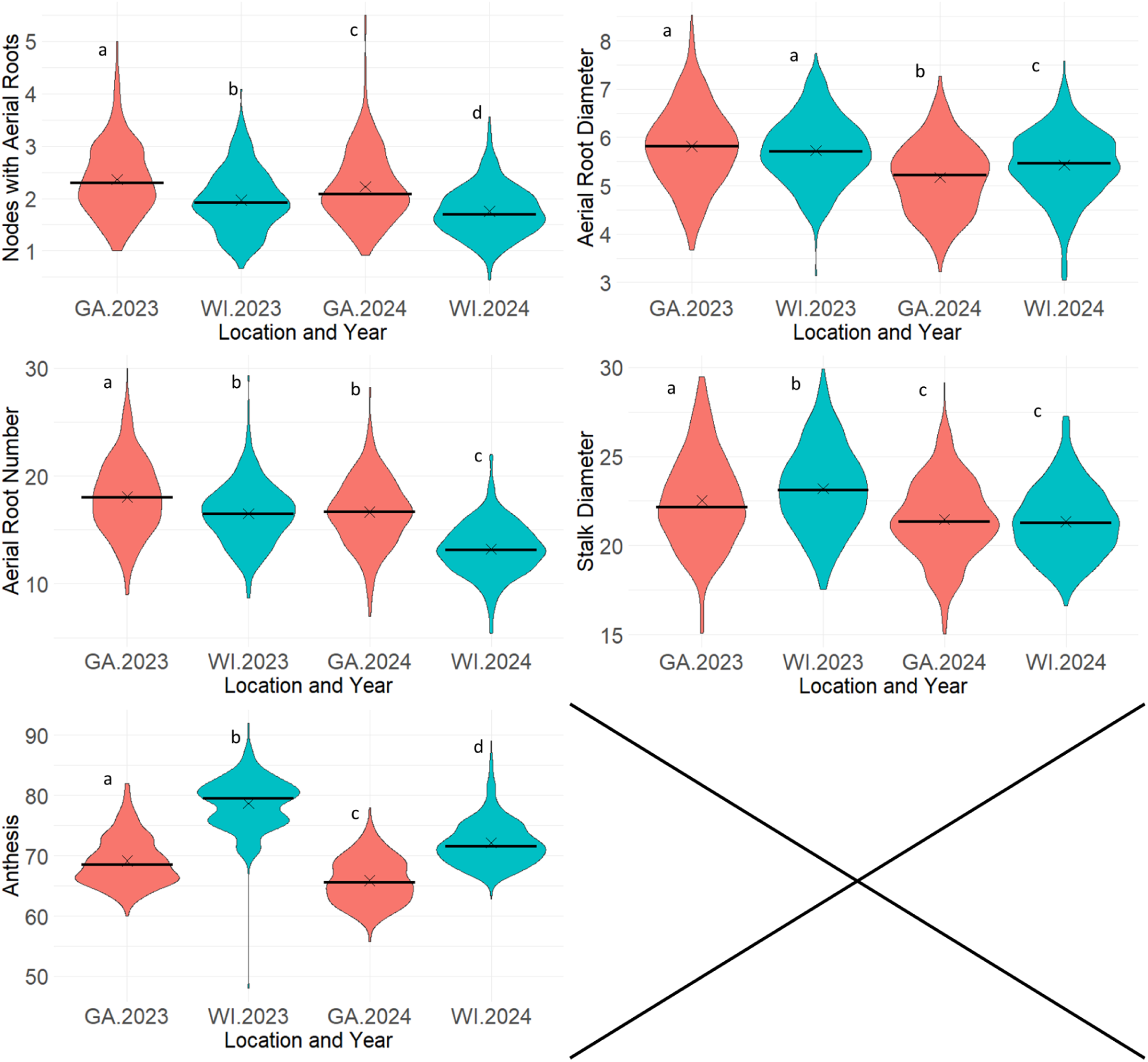
Violin plots of filtered phenotypes separated by year, and location. The mean for each group is marked with an “X” and the median is denoted by a black horizontal line. We observe significant differences between all years and locations in the Nodes with aerial roots phenotype. Aerial root diameter was found to be significantly higher in 2023 compared to 2024. The Wisconsin location had a significantly lower aerial root number in 2024 compared to the previous year. Stalk diameter was observed to be relatively consistent across locations and years. Anthesis was significantly earlier in the GA location. Differences between groups were identified with a Tukey test using the Agricolae R package.

**Figure S5.**
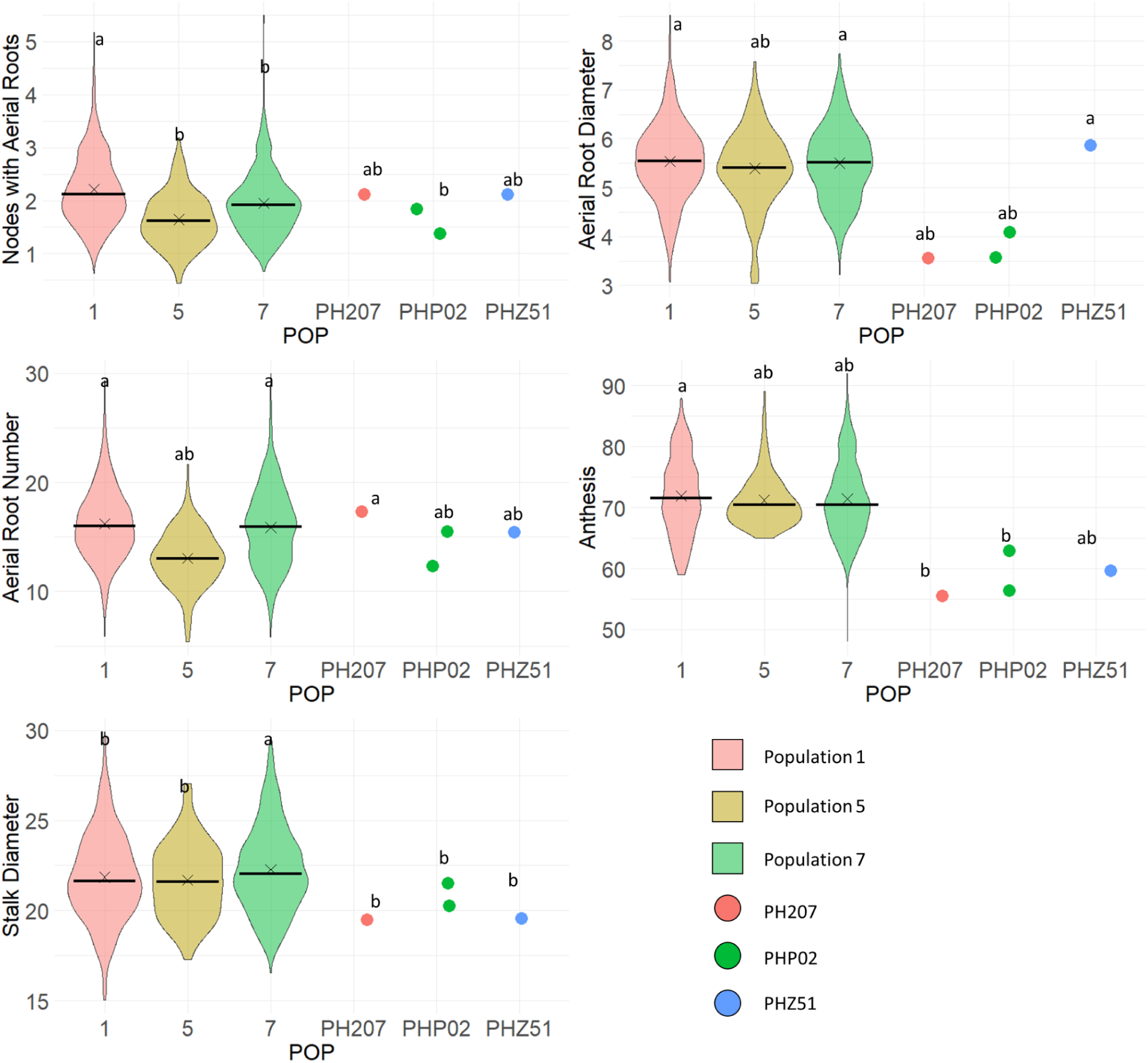
Violin plots of filtered phenotypes separated by population. The mean for each group is marked with an “X” and the median is denoted by a black horizontal line. Differences between groups were identified with a Tukey test using the Agricolae R package.

**Figure S6.**
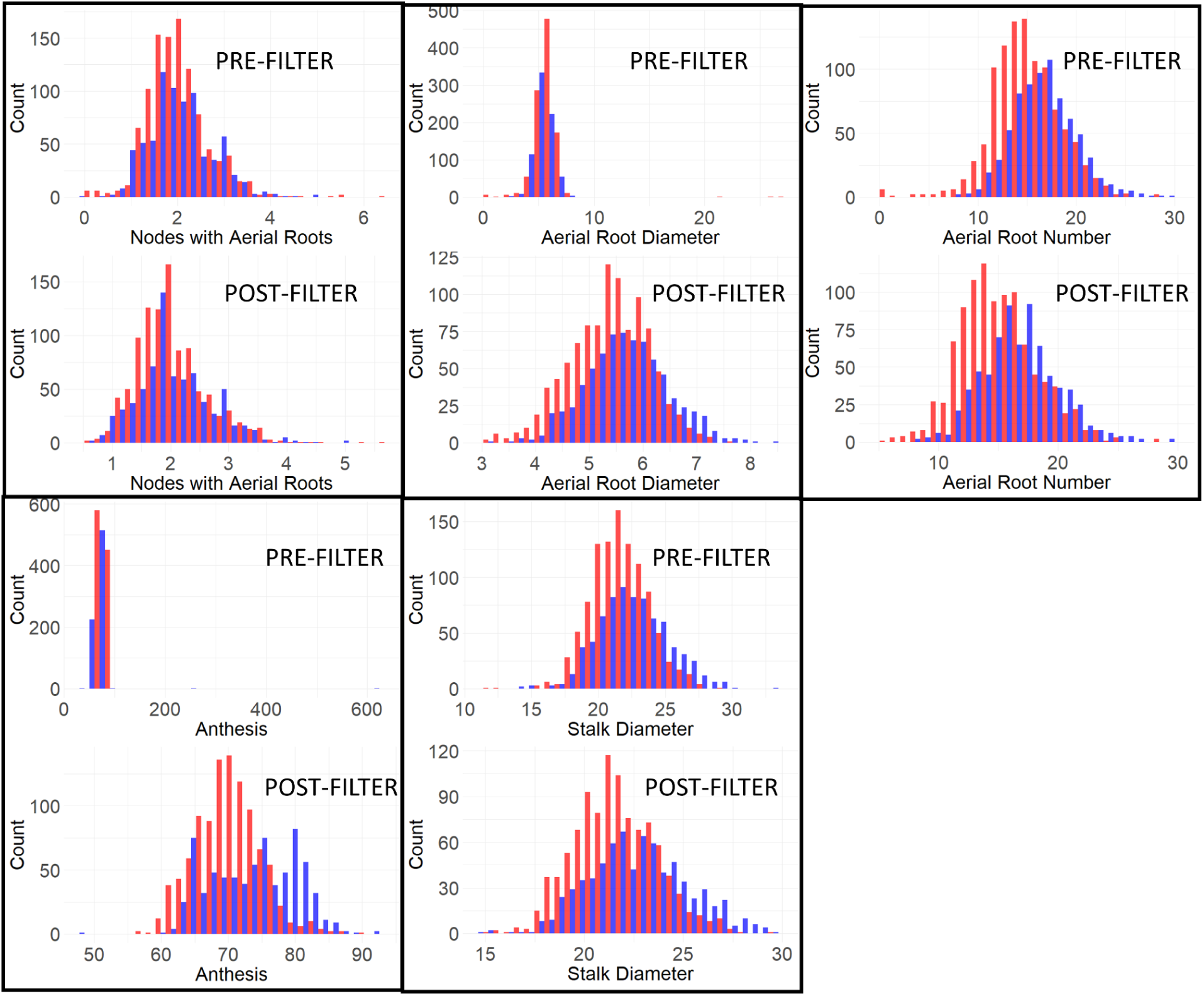
Distribution of Phenotypes in 2023 and 2024 for all measured populations (1, 5, and 7) pre and post filtering. Blue bars represent data from both GA and WI in 2023, and red represent data from both GA and WI in 2024. Phenotypes were filtered in R to remove erroneous outliers, keeping the data with the following parameters: nodes with aerial roots (>=0, <=6), aerial root diameter (>=3, <=10), aerial root number (>=5, <=30), Stalk Diameter (>=15, <= 30), and Anthesis (>=40, <= 100).

**Figure S7.**
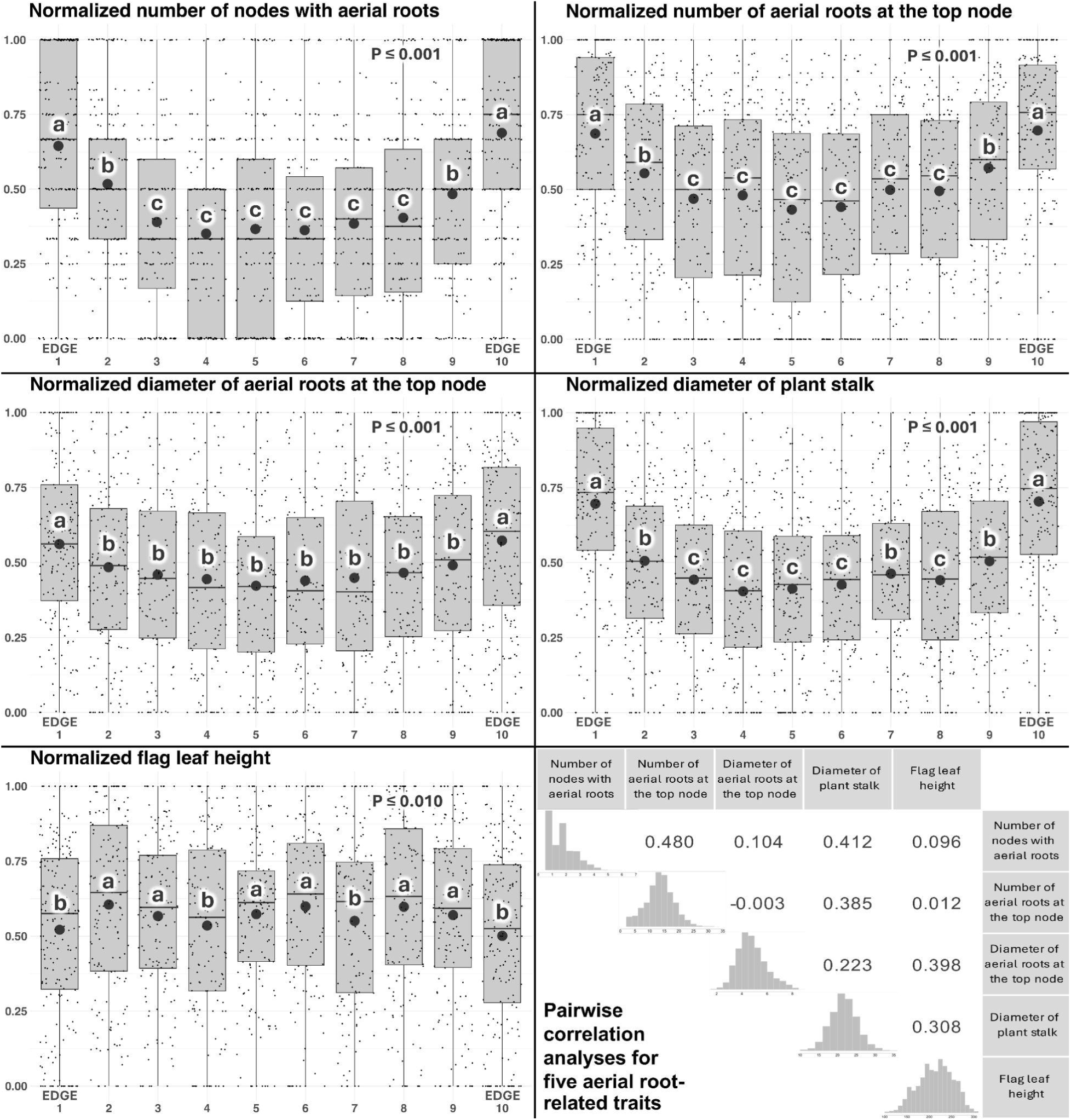
Field edge effects on five maize aerial root-related traits. On the x-axes of the five boxplots are the *deca*-percentile groups regarding plants’ position, calculated based on the position of each plant within the rows. Groups 1 and 10 constituted the row ends, so plants in these groups were the most exposed to edge effects. On the y-axes of the five boxplots are the normalized values observed for each trait in plants from different *deca*-percentile groups. The spheres represent the means of these values. The Scott-Knott clustering algorithm at 5% of significance level was applied for the mean comparison test. Lowercase letters (a, b, c) refer to the grouping of means. In the bottom right corner are the pairwise correlation analyses for those five traits, performed with the raw data, i.e., before they have been normalized on a 0 to 1 range.

**Figure S8.**
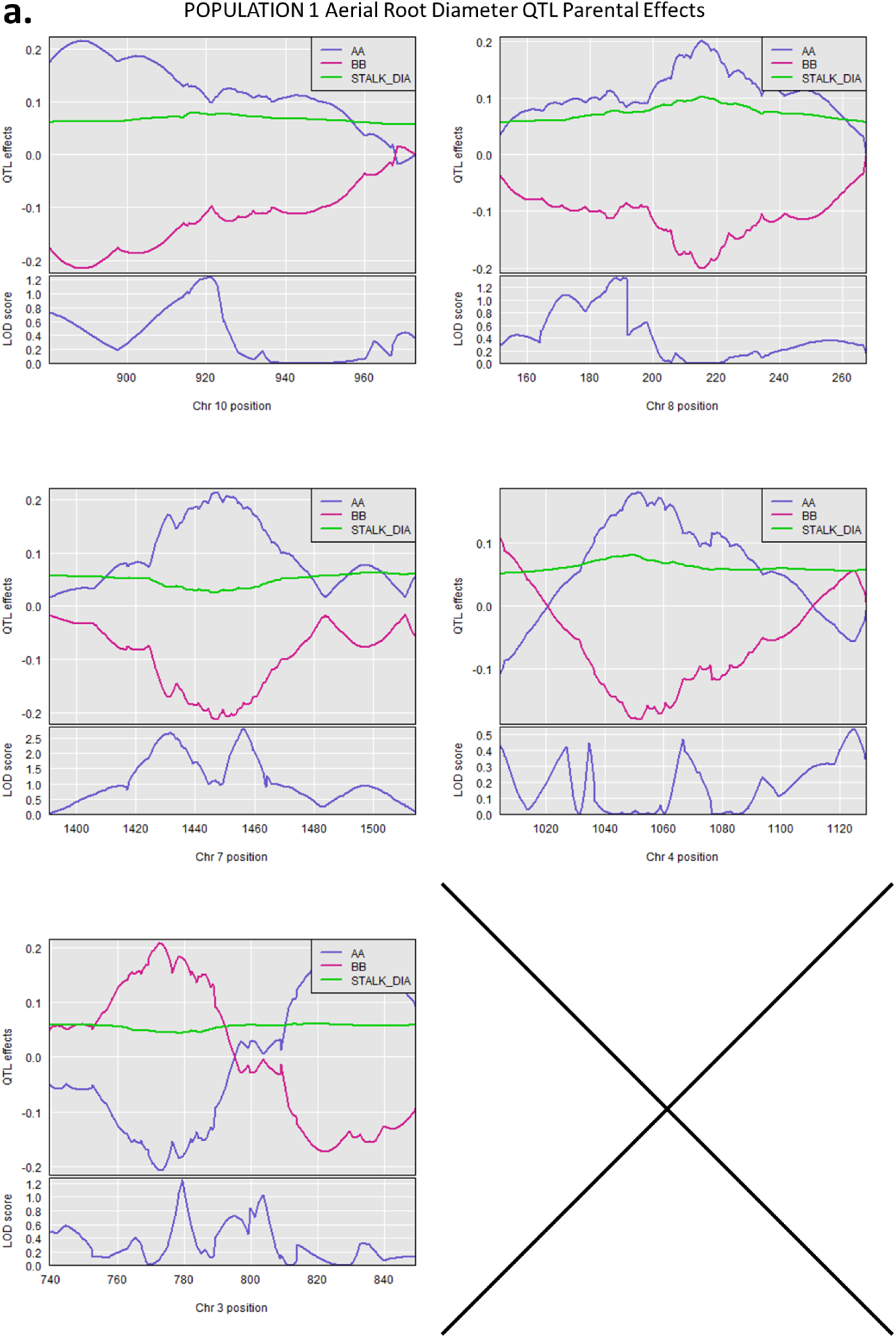
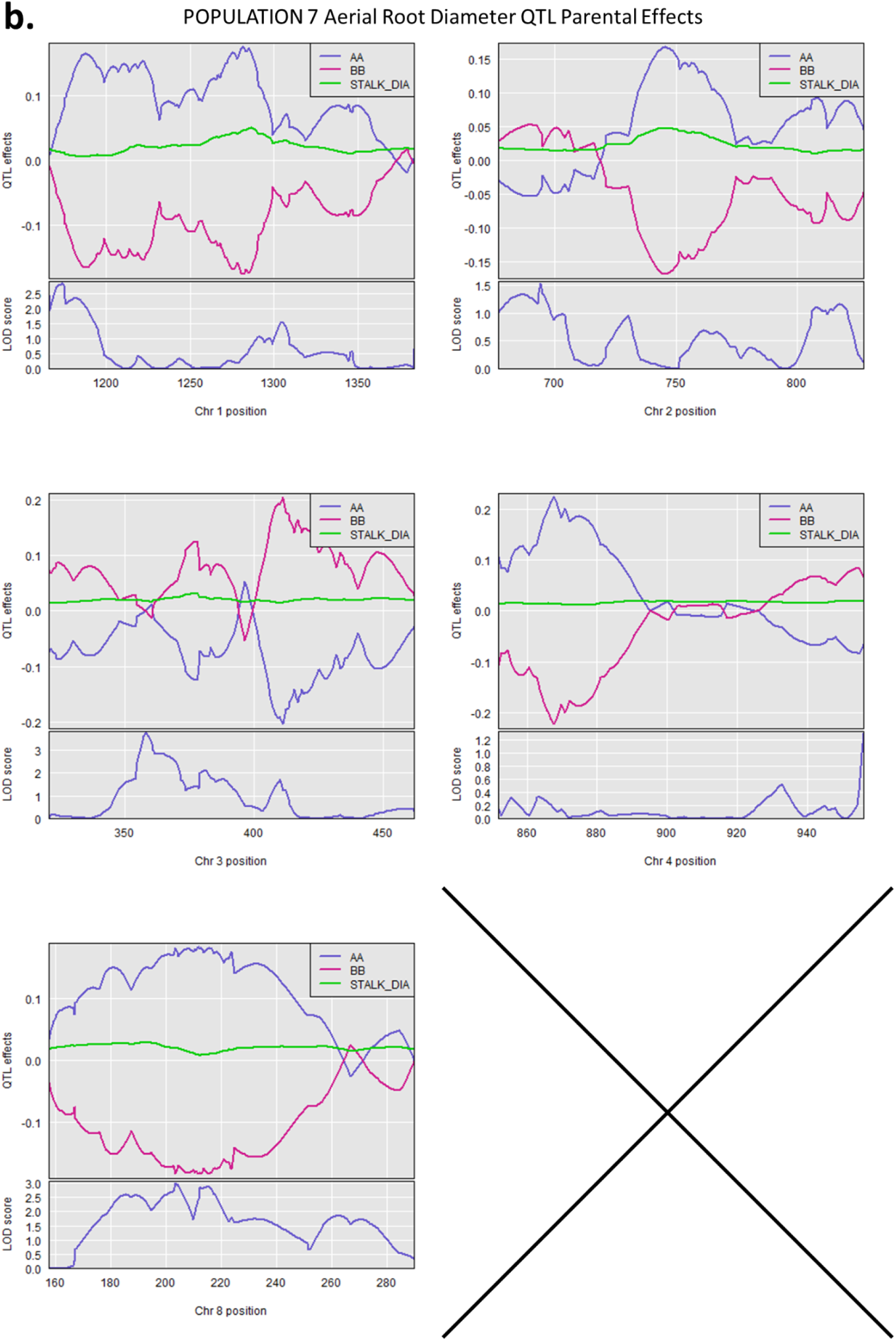
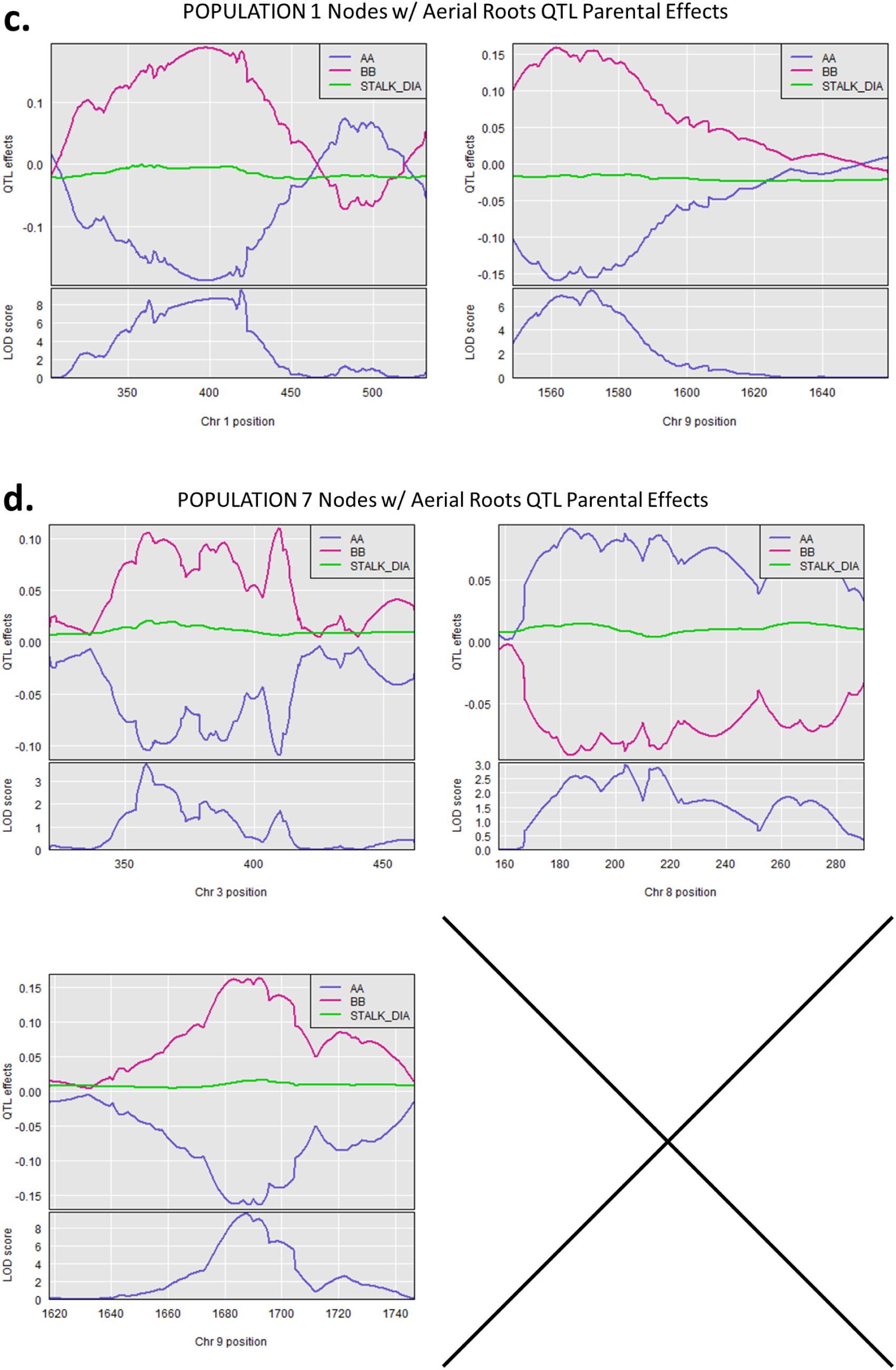
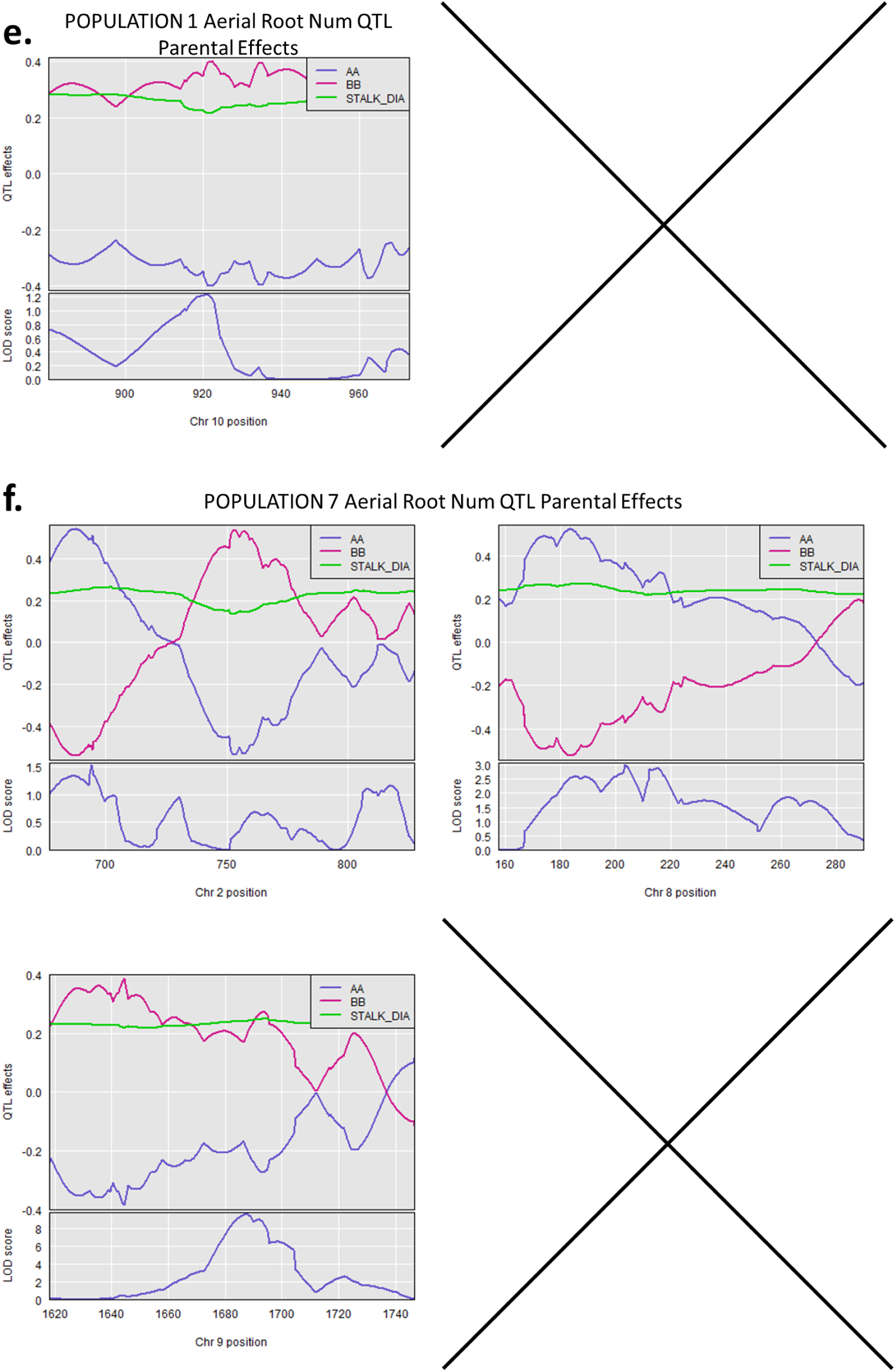
QTL parental effects for QTL in Populations 1 and 7. Aerial root diameter QTL for POP1 (a) and POP7 (b). Nodes with aerial roots QTL for POP1 (c) and POP7 (d). Aerial root number QTL for POP1 (e) and POP7 (f). AA genotype refers to the elite parent PHZ51 and BB corresponds to the landrace parent for POP1 and POP7. In general we observe positive effects to aerial root diameter originating from PHZ51 and positive effects to the nodes with aerial roots trait originating from the landrace parents. Note the LOD scores in these plots are derived from fitting a specific QTL model at a fixed position (e.g., a peak marker). These LOD scores reflect how well the model at that position explains the trait variation. As such they are independent of LOD scores generated for QTL across the genome seen in **Figure 5** of the main text.

**Figure S9.**
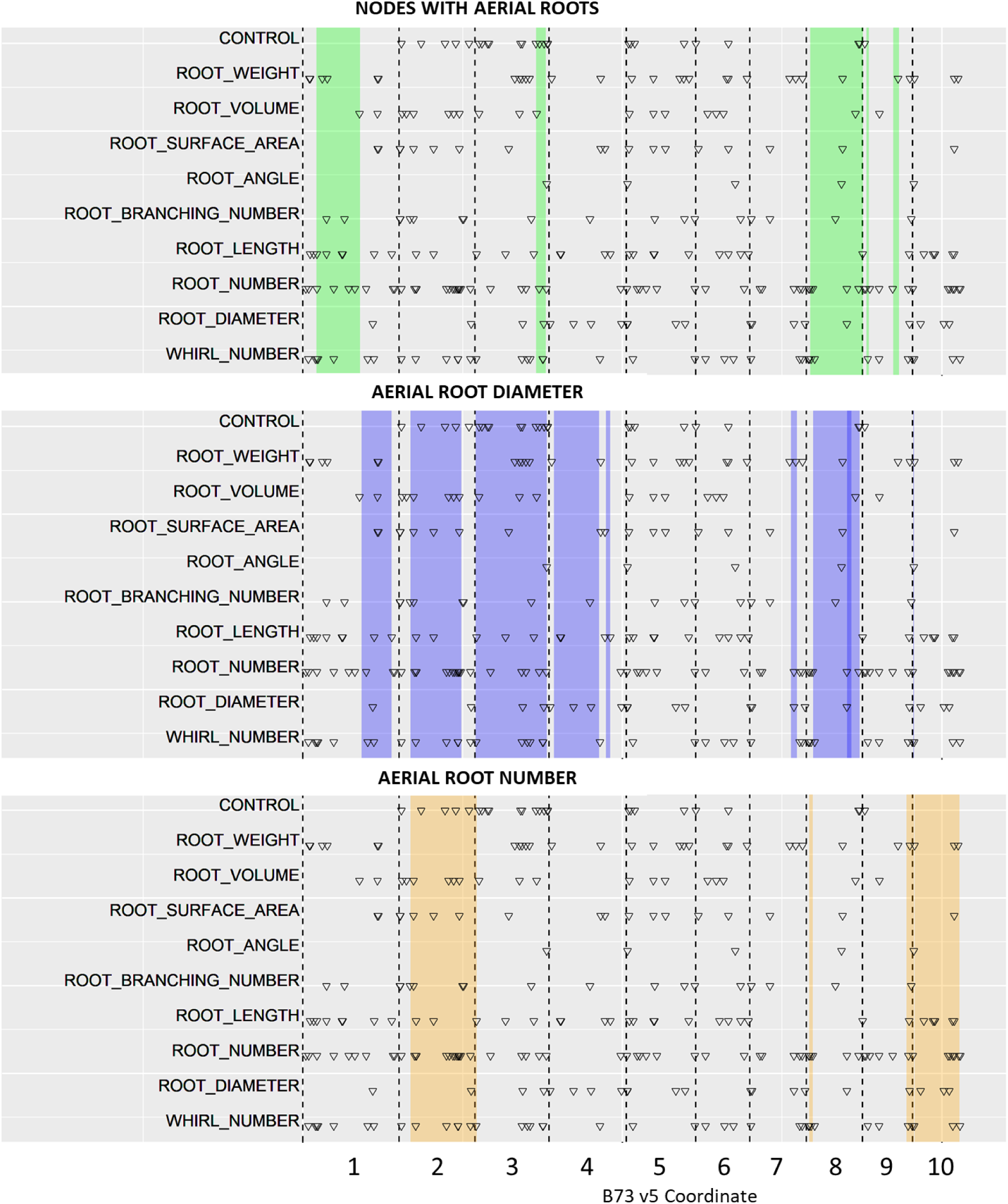
Overlap of published QTL/GWAS hits from 10 studies with QTL identified in populations 1 and 7. QTL positions are shown in the context of the B73 v5 genome. Shaded regions correspond to the positions of QTL identified in this study. Published QTL for 10 root trait classes are marked with black triangles.

**Figure S10.**
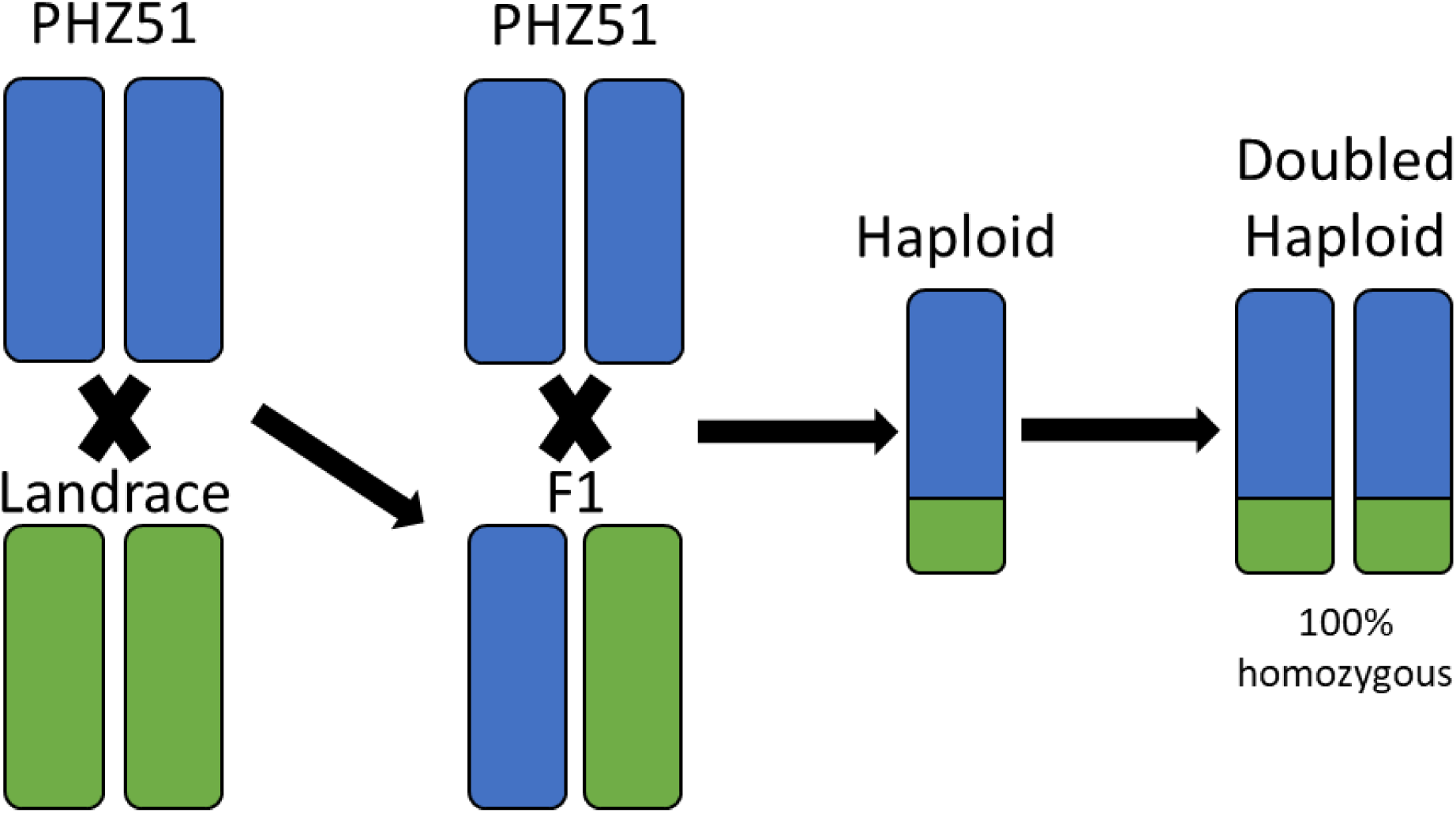
Crossing scheme for generating doubled haploid populations. Crosses were generated between one of 3 landraces (GRIN-PI-629238, GRIN-Ames-19897, GRIN-PI-645940) and the Ex-PVP line PHZ51 selected for its large diameter brace roots. The F1 lines were backcrossed to PHZ51 and the resulting BC1 lines. Limagrain conducted haploid induction and haploid doubling using the BC1 lines. Eight populations of doubled haploids were generated.

**Figure S11).**
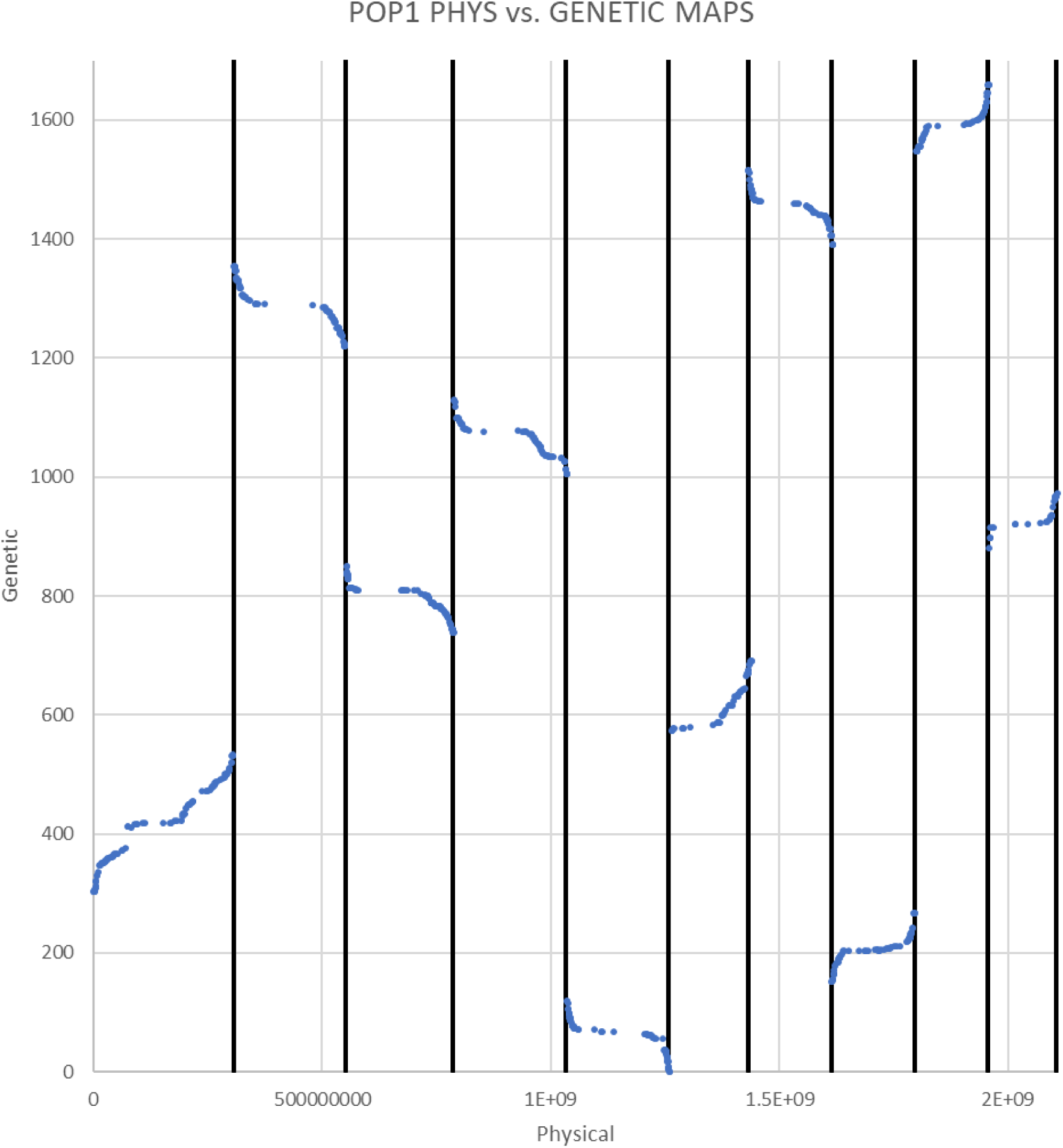
Comparison of physical and genetic maps for POP1. We do not observe any genetic markers that are located far from their physical position indicating our genetic map is accurate. The estimated genetic position for each marker is located on the Y axis. The X-axis represents the markers physical location in the B73 v5 genome.

**Figure S12.**
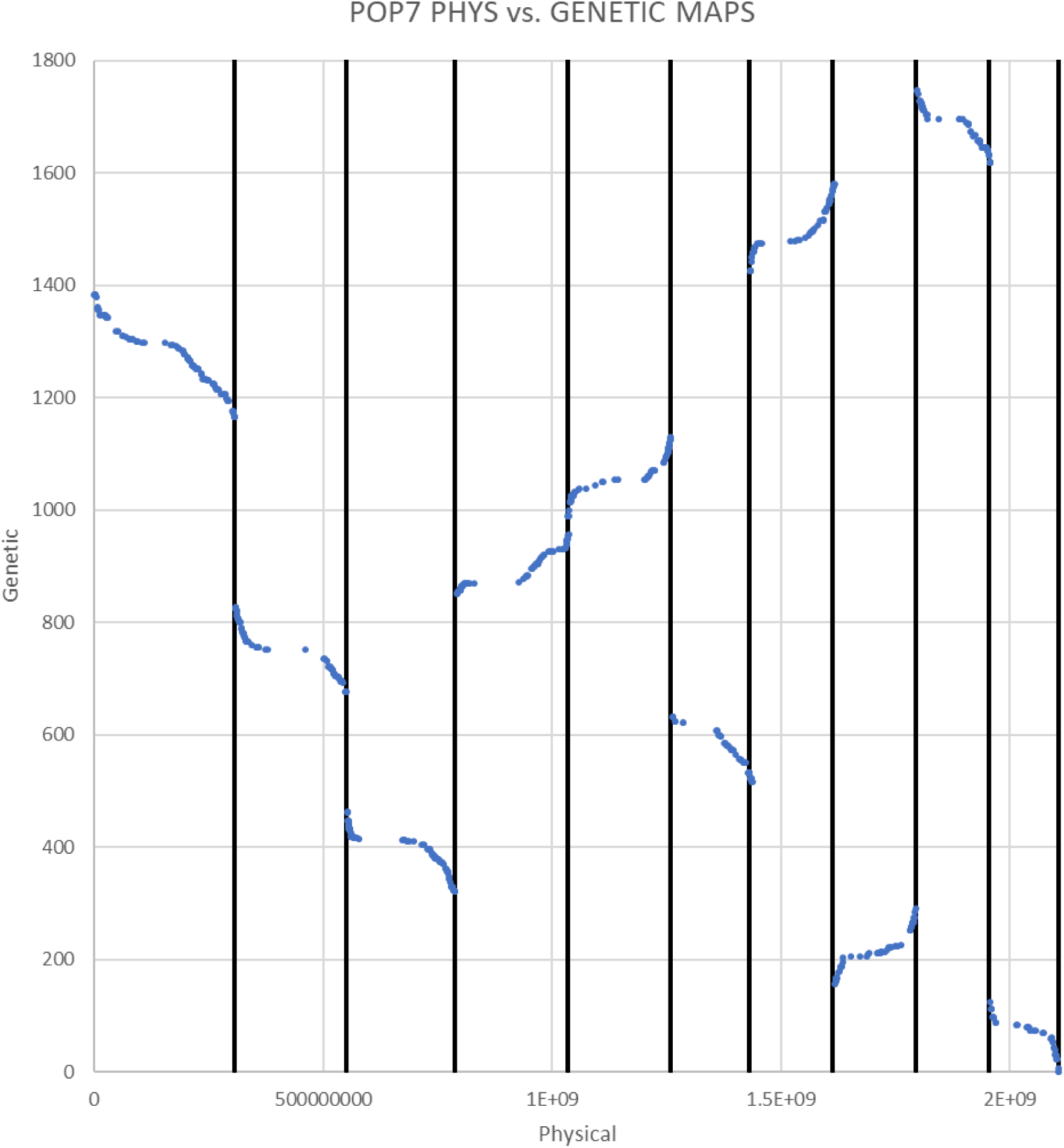
Comparison of physical and genetic maps for POP7. We do not observe any genetic markers that are located far from their physical position indicating our genetic map is accurate. The estimated genetic position for each marker is located on the Y axis. The X-axis represents the markers physical location in the B73 v5 genome.

## Supplemental Tables

**Table ST1.** nitrogen-fixing and non-fixing bacteria isolated from the rhizosphere of cereals and the mucilage of maize and sorghum. The strains were used for the following experiments: Acetylene Reduction Assay (ARA) and ^15^N isotope dilution assay (^15^N). The reference of the strains is provided below

| Diazotroph | Experiment | Reference | Source |
| --- | --- | --- | --- |
| *Azorhizobium caulinodans* ORS571<sup>T</sup> (Fixing strain) | ARA | (Dreyfus and Dommergues 1981) | Nodules in *Sesbania rostrata* |
| *Azospirillum baldaniorum* Sp 245<sup>T</sup> (Fixing strain) | ARA | (Baldani *et al.* 1983) | Wheat |
| *Azospirillum brasilense* FP2 (Fixing strain) | ARA | (Pedrosa and Yates 1984) | Wheat, maize and forage grass (*Poa pratense*) |
| *Azospirillum brasilense* FP10 (Non-fixing strain) | ARA | (Pedrosa and Yates 1984) | Wheat, maize and forage grass (*Poa pratense*) |
| *Azotobacter vinelandii* DJ (Fixing strain) | ARA | (Setubal *et al.* 2009) | Ubiquitous |
| *Azotobacter vinelandii* DJ100 $\Delta nifD$ (Non-fixing strain) | ARA and $^{15}\text{N}$ | (Robinson *et al.* 1986) | Ubiquitous |
| *Herbaspirillum seropedicae* SmR1 | ARA | (Pedrosa *et al.* 2011) | Maize, sorghum, rice, wheat and sugarcane |
| *Klebsiella michiganensis* A2 (Fixing strain) | ARA and $^{15}\text{N}$ | (Venado *et al.* 2025) | Maize mucilage |
| *Klebsiella michiganensis* A2 $\Delta nifH$ (Non-fixing strain) | $^{15}\text{N}$ | This study | Maize mucilage |
| *Klebsiella variicola* A3 (Fixing strain) | ARA | (Venado *et al.* 2025) | Sorghum mucilage |
| *Pseudomonas stutzeri* A1501 (Fixing strain) | Fixer strain | (Qiu *et al.* 1981) | Rice |
| *Paraburkholderia silvatlantica* SRMrh-20<sup>T</sup> (Fixing strain) | Fixer strain | (Perin *et al.* 2006) | Maize |

**Table ST2.**
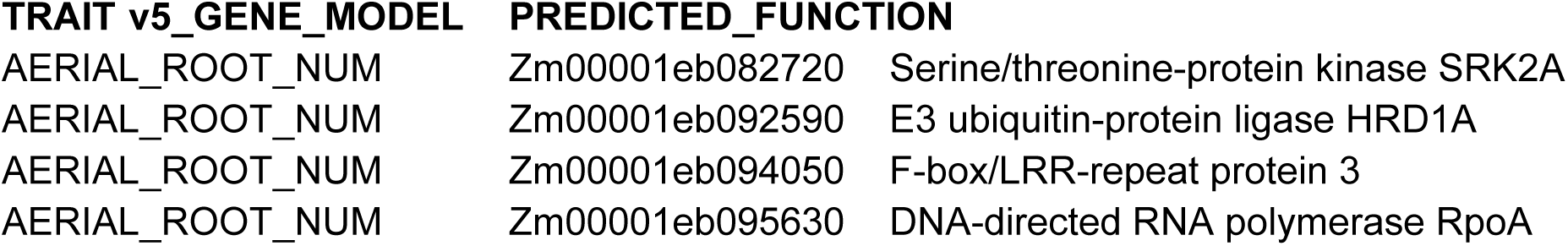
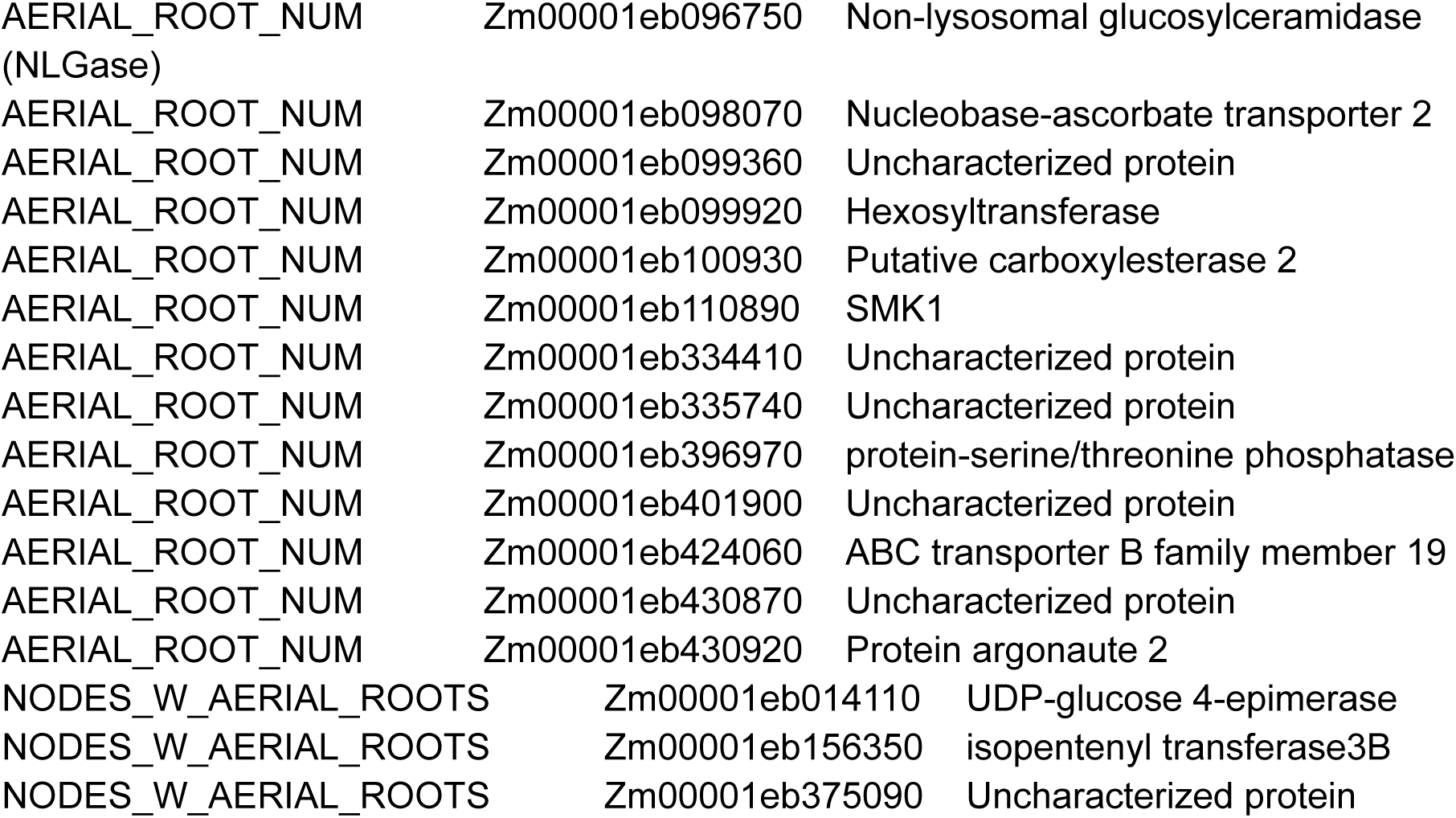
Putative candidate genes. Candidate gene modes from the B73 v5 annotations that fall within one of our QTL, and are associated with a published GWAS hit for the same trait. Note that we cannot be certain the same nucleotide change is present in our populations (limited by marker density).

